# Insights into adhesive and neuronal cell populations of the chaetognath *Spadella cephaloptera* using a single-nuclei transcriptomic atlas and genomic resources

**DOI:** 10.1101/2025.01.31.635879

**Authors:** Cristian Camilo Barrera Grijalba, June F Ordoñez, Juan Montenegro, Tim Wollesen

**Author notes:** To whom correspondence should be addressed: Tim Wollesen, Cristian Camilo Barrera Grijalba.

## Abstract

To cope with extreme environmental conditions diverse marine species have developed mechanisms that allow them to permanently or temporarily attach to substrates. In the intertidal zone of marine habitats, where tidal ranges and currents may drift organisms away from their habitat, temporary adhesive systems such as the one inherent the arrow worm *Spadella cephaloptera* (Chaetognatha) constitute an essential trait for the survival of this taxon. The underlying molecular mechanism of this system has not been described yet, and the existing morphological information is limited to adults. Furthermore, a relationship between the nervous system and the attachment in *S. cephaloptera* remains to be demonstrated. In this study, single-nuclei sequencing of *S. cephaloptera* hatchlings was performed, using as a reference a newly sequenced and assembled genome to identify the transcriptomic profiles of the cells mediating attachment, neuronal populations, and the main cell types of chaetognath hatchlings. Our findings, supported by previous studies, suggest that the chaetognath adhesive system evolved convergently to those of other other metazoans. Moreover, diverse cell types were identified in the ventral nerve center and multiple ciliated cell types previously described from anatomical observations were validated. Ongoing in-depth investigation of these data, together with datasets from other developmental stages, will provide further insights into the evolutionary origins of the unique chaetognath body plan.

## Introduction

Marine invertebrates rely on attachment for feeding, locomotion, and defense [1]. In this sense, there is a high diversity of clade-specific mechanisms regarding the attachment process [2], and these systems may be present across different developmental stages (e.g., hatchlings, and adults), exhibiting different stage-specific features [3][4]. During ontogeny, the cells associated with attachment can shift their transcriptomic program, resulting in both functional and location changes in the organism [4]. In marine invertebrates, the attachment systems have been categorized by features, such as duration (temporary vs. permanent) and structure (single-vs. multi-unit organs) [2]. Temporary attachment provides reversibility, enabling the organisms to interact more dynamically with their environment [1]. Such systems may feature specialized structures, such as multiple feet, which increase the contact area with the substrate, and mediate a stronger attachment [1].

Furthermore, it has been demonstrated that some taxa also secrete “glues” primarily composed of proteins, glycans, and lipids [5][6]. In this regard, secreted proteins mediating attachment in marine invertebrates often share conserved protein architectures [2][5]. Remarkably, in the case of non-permanent attachment, it has been suggested that the similarity in the amino-acid profile of those proteins results from convergent evolution shaping these mechanisms [2].

By contrast, permanent attachment typically occurs at a specific developmental point in the life cycle of marine invertebrates, often transitioning from a free-swimming larva to a sessile adult. [5][6]. This strategy allows the organisms to fix themselves to suitable locations using attachment organs that could be either single-unit or multiple units, and it is further mediated by a secretion composed almost entirely of cement proteins that curate over time enabling firm anchorage [2]. Remarkably, it has been observed that for the studied representatives from different taxa, the degree of similarity in the amino-acidic composition of the proteins involved in permanent attachment is lower than that observed in non-permanent attachment systems [2]. In addition, research on barnacles, which provide a classic example of permanent adhesion, underscores the role of attachment during ontogeny. During early development, the barnacle larvae (cyprid) benefit from temporary adhesion to explore before releasing permanent adhesive and continuing development [4]. Hence, understanding the molecular mechanisms associated with adhesive systems, whether temporary or permanent, could also offer insights into the broader developmental programs of marine invertebrates.

Despite extensive characterization of attachment processes in some phyla, a number of marine spiralian groups, (e.g., chaetognaths) remain significantly unexplored regarding the molecular mechanisms underlying their attachment system [5]. In this sense, chaetognaths have recently been considered to belong to the Gnathifera, the sister clade of the spiralian Lophotrochozoa (Fig. 1A) [7]. However, their phylogenetic position has been and is still debated [8]. In this view, the research on the adhesive system of benthic chaetognaths could therefore unveil molecular mechanisms for attachment while expanding the understanding of marine invertebrate evolution.

**Figure 1.**
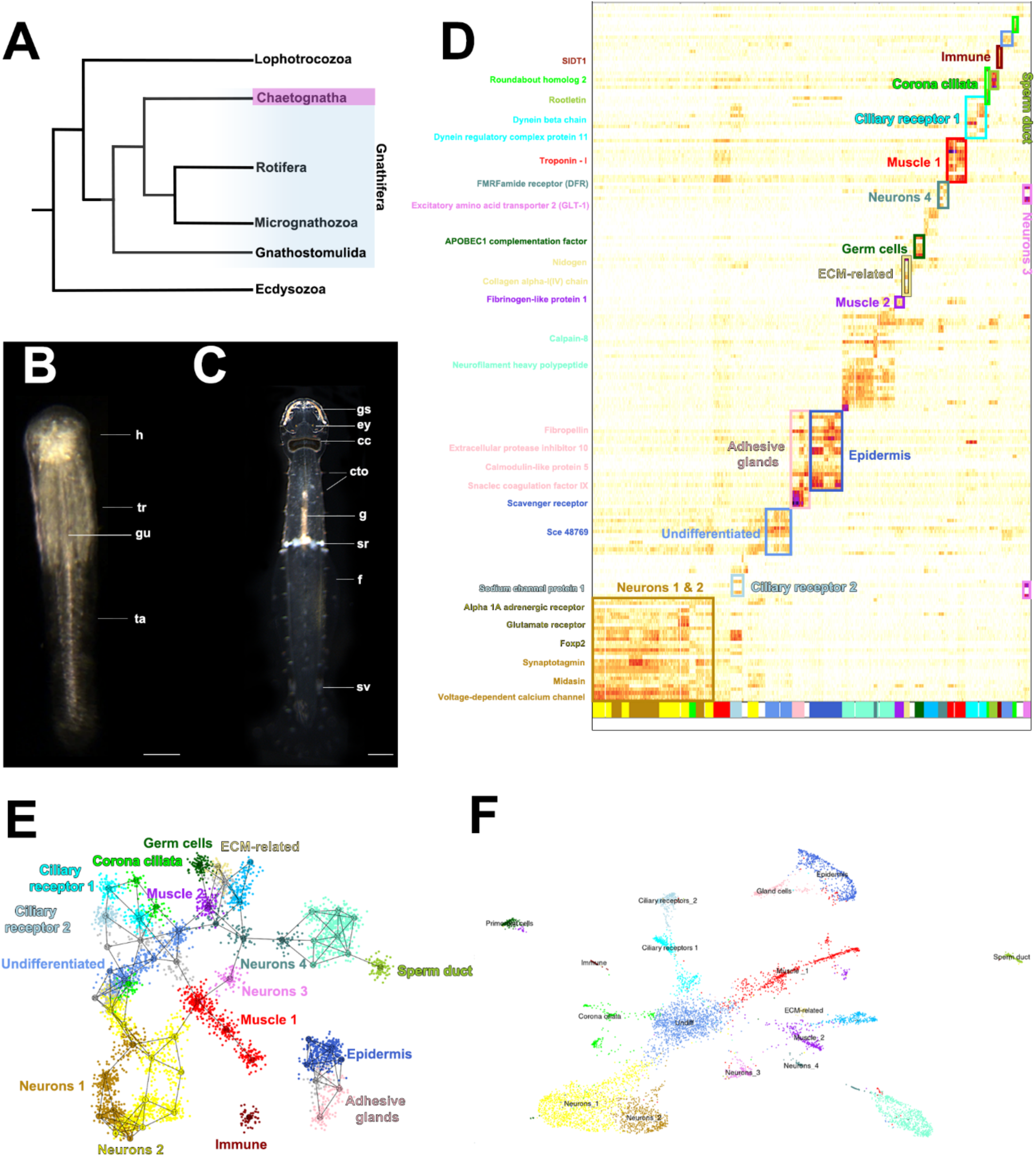
Single nuclei transcriptomics in the hatchling of *S. cephaloptera* identify an adhesive cell gland population along different main cell populations. **A.** Phylogenetic placement of *S. cephaloptera* within protostomes, illustrating its position within the Gnathifera clade (phylogeny after [7]. **B.** Dark-field microscopy of the hatchling stage of *S. cephaloptera*, indicating the three main body regions: head (h), trunk (tr), and tail (ta). **C.** Dark-field microscopy of the adult stage, depicting key anatomic features, including grasping spines (gs), and the corona ciliata (cc) in the head, ciliary tuft organs (cto) and seminal receptacles (sr) in the trunk, and seminal vesicles (sv) in the tail. **D.** Heatmap of the expression of 195 gene markers across the 57 identified metacells (most informative genes are depicted). **E.** Two-dimensional projection of the 57 metacells represented with numbers **F.** UMAP representation of the 18 identified cell clusters. Scale bar in C: 100 µM; Scale bar in D: 200 µM. Additional abbreviations: ey, eye; gu, gut; f, fin.

*S. cephaloptera*, the probably best studied benthic chaetognath (Fig. 1A, B), inhabits the intertidal zone, and its temporary attachment based on multiple adhesive papillae in the ventral surface, allows dynamic interactions with the environment and is evident from the hatchling stage onward [8][9]. Furthermore, the cells hypothesized to mediate the attachment process undergo changes in their distribution during ontogeny, suggesting an important role in development [10]. Notably, there is a hypothesis that these adhesive cells are innervated, although evidence is still lacking [8].

To date, descriptions of the different cell types of *S. cephaloptera* derive from Transmission Electron Microscopy (TEM) analyses in adults [8], [11], [12]. More recently, a hatchling-stage study examined the nervous system using a candidate-gene approach. However, the putative neuronal system related to the adhesive process has not been studied [13]. Meanwhile, the use of single-cell sequencing has enabled extensive transcriptomic profile diversity analyses in marine invertebrates, revealing the molecular basis underlying taxon-specific cell types [14][15][16][17]. Despite the utility of this approach, single-cell or single-nuclei transcriptomics remain unexplored in chaetognaths.

In the present study, the integration of genomic, transcriptomic, and single-nuclei sequencing data sets allowed to characterize genes associated with the putative adhesive cells in hatchlings of *S. cephaloptera*. Furthermore, by comparing these findings with data from other marine invertebrates, the results provide support for the underlying convergent evolution hypothesis of attachment processes.

## Methods

### Animal collection and fixation

Adult specimens of *S. cephaloptera* were collected along the shore near the Station Biologique de Roscoff in Roscoff, France, and transferred to petri dishes to induce reproduction and egg-laying. Eggs were collected in separate dishes to monitor hatching times. After 8-24 hours (1dph) specimens intended for *in-situ* hybridization (ISH) were relaxed with MgCl2 and fixed in 4% paraformaldehyde (PFA), prepared in 0.5 M MOPS, 5 mM EGTA, 10 mM MgSO4, 2.5M NaCl, at 4°C overnight. Fixed specimens were then washed thrice for 10 min in pre-chilled methanol and stored at –20 °C. For RNA-seq and single-nuclei sequencing experiments, the specimens were flash-frozen in low-binding tubes using liquid nitrogen.

### Genome sequencing and assembly

High molecular weight (HMW) DNA was extracted from a single adult specimen of *S. cephaloptera* using the HMW Monarch Kit (New England BioLabs). Pac-Bio HiFi libraries were prepared using the SMRTbell Express Kit and sequenced on a Sequel II system by the Next Generation Sequencing Facility at Vienna BioCenter Core Facilities (VBCF) in Vienna, Austria. The obtained reads were filtered for adapter sequences using HiFiAdapterFilt [18] and subsequently assembled using Hifiasm (0.19.8) [19], passing an estimated genome size of (800 Mbp), determined by Feulgen densitometry. Assembly quality was assessed using QUAST (5.2.0) [20], and the completeness was evaluated using BUSCO (5.8.2) [21] with the metazoa_odb10 database. The best haplotype (h1) based on BUSCO and QUAST stats was selected for downstream analyses. Contiguity of the assembly was improved using P_RNA scaffolder (https://github.com/CAFS-bioinformatics/P_RNA_scaffolder) with bulk RNA-seq Illumina reads [22]. Repetitive regions were identified *de-novo* using RepeatModeler2 [23], and soft-masked with RepeatMasker [24]. Genome gene structure annotation was performed with the BRAKER3 (v3.0.8) pipeline [25–37], integrating RNA-seq hints from *S. cephaloptera* bulk Illumina reads [22] mapped to the genome using HISAT2 (2.1) [38]. Additionally, protein hints were provided by means of the metazoan protein database from OrthoDB v.11 [39].

### Transcriptome sequencing and assembly

Total RNA was extracted from flash-frozen individuals of *S. cephaloptera* representing four developmental stages eggs, 24 hours post-hatching, 5 days post-hatching, and adults) using the RNAqueous Micro Total RNA Isolation kit (Invitrogen). The RNA elutions were pooled in a 5-3-2-2 ratio, respectively. Library preparation was carried out using the Pac-Bio Kinnex full-length RNA kit, followed by sequencing on a Revio system. The obtained reads were processed according to the Isoseq3 (v4.0.0) pipeline (https://github.com/PacificBiosciences/IsoSeq). In brief, HiFi reads were segmented using *skera*. Then, after primer removal and refinement the isoforms were clustered using *cluster2.* The isoforms were then mapped to the reference genome using *pbmm2* and finally collapsed with the *collapse* function. Resulting isoforms were further clustered by CD-HIT (V4.8.1) [40], and proteome completeness was assessed using BUSCO (5.8.2) [21].

### Functional annotation

Gene transfer format (GTF) files from Braker3 and Isoseq3 were merged using in-house scripts, prioritizing in all cases the evidence provided by the Iso-Seq data. The resulting merged GTF was used to extract the transcript sequences, which were processed with TransDecoder (v5.7.1) [41]. TransDecoderPredict was guided on supported blastp hits to improve accuracy (NCBI blastp v2.15.0 against the SWISS-PROT database) [42] [43]. The predicted coding transcripts were functionally annotated using HMMER-PFAM 3.4 [44] and eggNOG-mapper (v2.1.12) [45].

### Single-nuclei data generation

Nuclei extraction was performed as described in [46] with modifications. Frozen hatchlings of *S. cephaloptera* were homogenized on ice in homogenization buffer (HB) (250mM sucrose, 25 mM KCl, 5mM MgCl2, 10mM Tris-HCL, pH = 8, 0.1% IGEPAL, 1uM DTT, 0.4 U/ µL RNAse Inhibitor (NEB), 0.2 U/µL Superasin (ThermoFischer Scientific), and allowed to stand for 5 min. Unlysed tissue was removed by low centrifugation, 100g for 1 min at 4°C. Nuclei were then pelleted at 400g for 4 min at 4°C and subjected to two additional HB washes/pelleting steps. The final nuclei were resuspended in Nuclei Storage Buffer (NSB) 430mM sucrose, 70 mM KCl, 2mM MgCl2, 100mM Tris-HCL, pH = 8, 0.4 U/µL RNAse Inhibitor (NEB), 0.2 U/µL Superasin (ThermoFischer Scientific), and counted using DAPI on a hemocytometer.

Extracted nuclei (n=20,000) were diluted in Dulbecco’s PBS (Sigma-Aldrich) and subsequently used for library preparation using Chromium Next GEM Single-Cell 3’ GEM, library & Gel Bead kits v3.1 (10X Genomics), and sequenced using Illumina NovaSeq SP platform. Reads were mapped to the reference genome using 10x Genomics CellRanger v7.1.0 and the merged GTF file.

### Single-nuclei data processing

The filtered count matrix from CellRanger was processed using Seurat (v4.4.2) [47]. Multiplets were removed based on the number of features and RNA count distributions. The filtered data were then normalized by applying a scale factor of 5000, selecting 2000 variable genes via the FindVariableFeatures function before scaling.

The number of dimensions for the UMAP and FindNeighbors was determined by PCA, selecting the minimum number of principal components that captured >90% of the variance. Cells were grouped using the function FindNeighbors, with the cosine distance and k=10. Clusters of cells were generated using FindClusters with a resolution of 0.42, producing 18 clusters. Marker genes for each cluster were identified using FindAllMarkers with a log-fold change threshold of 1, selecting for the markers that are upregulated and present in at least 10% of cells in a cluster. The distribution of these markers guided the putative identity assignments to the different clusters.

In parallel, the filtered matrix was analyzed using the Metacell R package [48]. Cells with fewer than 1000 UMIs were filtered out based on the UMI count distribution. Then, genes with a scaled variance of ≥0.08 and ≥ 100 UMIS in the dataset were selected. This subset of genes was used to generate a similarity graph (k =100). After resampling and calculating the co-clustering graph, outlier cells were removed from each metacell based on their expression profile. Marker genes were then colored based on their functional annotation, and their distribution was reviewed in a heatmap and a 2D projection to assign the identities of metacells. Finally, relationships between individual metacells were evaluated using a confusion matrix.

### Riboprobe synthesis and whole-mount fluorescent *in situ* hybridization

Multiple developmental stages of flash-frozen individuals of *S. cephaloptera* were used for total RNA extraction using the RNAqueous Micro Total RNA Isolation kit (Invitrogen). Then, a cDNA template was generated using the First Strand cDNA Synthesis Kit for RT-PCR (Roche).

From selected cell-type markers, PCR primers were designed using Primer-Blast with an annealing temperature of 60°C. The reverse primer included a T7 promoter sequence at the 5’ end (TAATACGACTCACTATAGGG). PCR amplicons were purified with a QIAquick PCR Purification Kit (Qiagen) and verified by Sanger sequencing. Purified products were then used as templates for riboprobe synthesis using DIG RNA labeling mix (Roche) and T7 polymerase.

For whole-mount fluorescent *in situ* hybridization (FISH) assays, all incubations were carried out at room temperature (RT) with gentle agitation (40 rpm), unless otherwise specified. Specimens were first incubated in *H*_2_*O*_2_ for 30 min in the dark, then rehydrated stepwise into PBT (1x PBS + 0.1% Tween-20) and washed 3x 5 min in PBT. Permeabilization was performed by incubating the samples in Proteinase-K (2µg/mL) in PBT at 37°C for 10 min. The reaction was stopped using ice-cold glycine (2mg/mL) followed by washes of 5 min in PBT. Samples were then transferred to PBS +1% triethanolamine and subsequently incubated in PBS+ 1% triethanolamine with 0.3% acetic anhydride for four 5 min steps. Post-fixation was performed in 4% PFA in PBT for 45 min, followed by four 5 min washes with PBT. Specimens were prehybridized in hybridization buffer (50% formamide, 5× SSC, 100 µg/mL heparin, 5 mM EDTA, 100 µg/mL yeast tRNA, 0.1% Tween 20, 5% dextran sulfate) for 16h at 62°C, and hybridization was carried out at 61.5°C in the same buffer with 2 µg/mL of riboprobe for 18h. To remove nonspecific probe binding, the samples were washed twice with hybridization buffer for 30 min, followed by 20 min washes of increasing concentrations of 2xSSC at 61.5°C (25%/50%/75%/100%-relative to hybridization buffer), and three 20 min washes of 1xSSC at 61.5°C. The samples were blocked for 3 hours in blocking solution (Blocking Reagent (Roche) in maleic acid), then incubated overnight with a DIG-labeled AP-antibody (2:2500 in blocking solution) at 4°C. After 8x 5 min washes with PBT, the colorimetric reaction was visualized with 0.25 mg/mL Fast Blue BB and 0.25 mg/mL NAMP (Sigma) in SB8.2 (0.1 M Tris-HCl pH 8.2, containing 50 mM MgCl2, 100 mM NaCl, 0.1% Tween-20) [9]. After the expression pattern reached the desired intensity, the samples were washed twice in PBT for 5 min, post-fixed with 4% PFA in PBT for 1 hour, and washed four times for 5 min in PBT. Finally, specimens were incubated in DAPI (Sigma-Aldrich) overnight and cleared with 2,2′-thiodiethanol (Sigma-Aldrich) and mounted for imaging on a Leica TCS SP5 confocal microscope (Leica Microsystems).

### Code availability

The scripts for data preprocessing, genome and transcriptome assembly, and subsequent single-nuclei analyses of the hatchling dataset, will be available on GitHub at https://github.com/cristiancb94/Single_nuclei_hatchling. The computations detailed in this project were carried out using the Life Science Compute Cluster (LiSC) at the University of Vienna. When the work was conducted, LiSC ran on Oracle Linux Server v9.5 and employed SLURM [49] for task management. Several tools were packaged within conda environments, as noted individually for each script where applicable.

## Results

### Genome assembly and annotation

After filtering the Pac-Bio Hi-Fi reads for adapter sequences removal, 99.8% of the reads were retained, resulting in 5,321,461 reads. The Hifiasm assembly size is about 1.18GB distributed across 12,407 contigs, with an N50 of 170 kB. Coverage estimates indicated 49x in homozygous regions and 21x in heterozygous regions, yielding a 2.3 homozygous-to – heterozygous coverage ratio. Mapping the filtered Hi-fi reads to the selected haplotype (H1) achieved a 99.14% mapping rate. BUSCO completeness analysis revealed 82.1% complete BUSCOs (65.4% single copy, 16.7% duplicated) with 3.5% fragmented and 14.4& missing, suggesting moderate completeness with some degree of duplication. Repeat masking pipeline identified 14.99% of the assembly as LTR elements and 2.97% as DNA transposons. BRAKER3 annotation produced 16,405 gene models with 76.5% complete BUSCOs (56.8% single copy, 19.7% duplicated), 3.9% fragmented, and 19.6% missing when evaluating the protein-coding sequences.

### Transcriptome assembly and annotation

Pac-Bio Iso-seq yielded 9,014,449 raw reads, which were split into 62,638,435 S-reads (mean length: 1,347 bp) after adapters removal, indicating a concatenation factor of 6.94. After initial clustering, 55,234,357 isoforms were obtained, all of which mapped to the genome assembly. Collapsing these isoforms produced 60,781 unique loci that were used as gene models for retrieving their corresponding genomic sequences. Further collapsing of these loci using CD-HIT produced a subset of protein-coding sequences with 85.2% complete BUSCOs (58.7% single copy, 26.5% duplicated), with 4.1% fragmented and 10.7% missing.

### Reference annotation file

Merging the BRAKER3 and Iso-seq GTF files, prioritizing the Iso-seq isoform families as a reference, resulted in 61,280 gene models, with 499 predicted features contributed by BRAKER3. From these gene models, 196574 transcripts were identified by TransdeCoder as coding sequences, and 12156 unique gene models were functionally annotated.

### Cell ranger

Using the Cell Ranger pipeline to map RNA-seq reads against the reference genome identified 7836; with an average depth of 56,278 reads per cell and a median of 742 genes detected per cell. The estimated fraction of reads inside cells was 27,6%, and the median UMI count per cell was 1,052. The mapping rate to the genome was 54.1%, with 31.3% of the reads confidently mapped to the transcriptome.

### Single-nuclei-analysis

Applying the Seurat pipeline unveiled 18 distinct cell clusters of cell populations (Fig. 1F). Four clusters corresponded to neuronal diversity: neurons 1 (1,284 cells), neurons 2 (540 cells), neurons 3 (143 cells), and neurons 4 (106 cells). Two additional cell populations were characterized by ciliary receptor genes: ciliary receptors 1 (380 cells) and ciliary receptor 2 (220 cells), plus one cell cluster related to the sensory organ Corona Ciliata (250 cells). Two muscle cell populations were identified: muscle 1 (657 cells) and muscle 2 (333 cells). Regarding the outer layer of cells in the hatchling, one epidermis cell cluster (543 cells) and one adhesive gland cell cluster (194 cells) were identified. One additional cluster was associated with primordial cells. Additional putative clusters include an extracellular matrix cell-associated cluster (74 cells), and a putative immune cell population (68 cells). In addition, one cluster of undifferentiated cells was also detected (1284 cells). Furthermore, using the Seurat pipeline allowed the identification of a comprehensive list of cluster-associated markers. Retrieved markers with their respective annotations are provided in Supplementary Table 1. This dataset is the basis for subsequent analyses and discussion, offering a comprehensive overview of the most significantly upregulated genes within each cluster. All the genes that were used for ISH are highlighted in green.

In parallel, the Metacell algorithm allowed the identification of 57 metacells and their relationships (Fig. 1D, E). Analyzing their identities revealed clusters consistent with those from Seurat, further corroborating the number of cell populations recovered by the single-nuclei extraction.

### Cluster validation

#### Neurons

After hatching, individuals of *S. cephaloptera* possess a nervous system that includes a cephalic cerebral ganglion and the ventral nerve center (VNC), which is composed of a pair of condensed lateral and medioventral neuronal somata clusters located ventrally in the trunk region [10]. Both Metacell and Seurat pipelines indicated four neuronal clusters, with the clusters neurons 1 and neurons 2 displaying closely related transcriptomic profiles. Nevertheless, they exhibit differentially expressed markers which hinds towards two distinct cell populations.

Cell cluster neurons 1 co-expresses adrenergic and acetylcholine receptors in ∼20% of the cells, whereas neurons 2 features a 3% higher proportion of cells expressing the neuronal marker *synaptotagmin-1* (*syt1*) compared to neurons 1 [14]. Consistent with this observation, the UMAP projection (Fig. 2A) localizes *syt1^+^* cells predominantly within neurons 2. On the other hand, FISH revealed *syt1^+^* cells lining the inner portion of both lateral somata clusters of the ventral nerve center along the sagittal axis (Fig. 2B, C). Additionally, the medio-posterior area of the trunk shows expression in two domains associated with bundles of neuronal nuclei that, in adults, give rise to bilateral neurons [8] (Fig. 2C, arrowheads). In the head region, ventrally, there are two additional bilateral expression domains in the proximity of the forming esophageal ganglia extending towards the cerebral ganglion (Fig. 2B, encircled stippled lines). This structure is not fully developed at this stage, which may contribute to the observed reduced expression.

**Figure 2.**
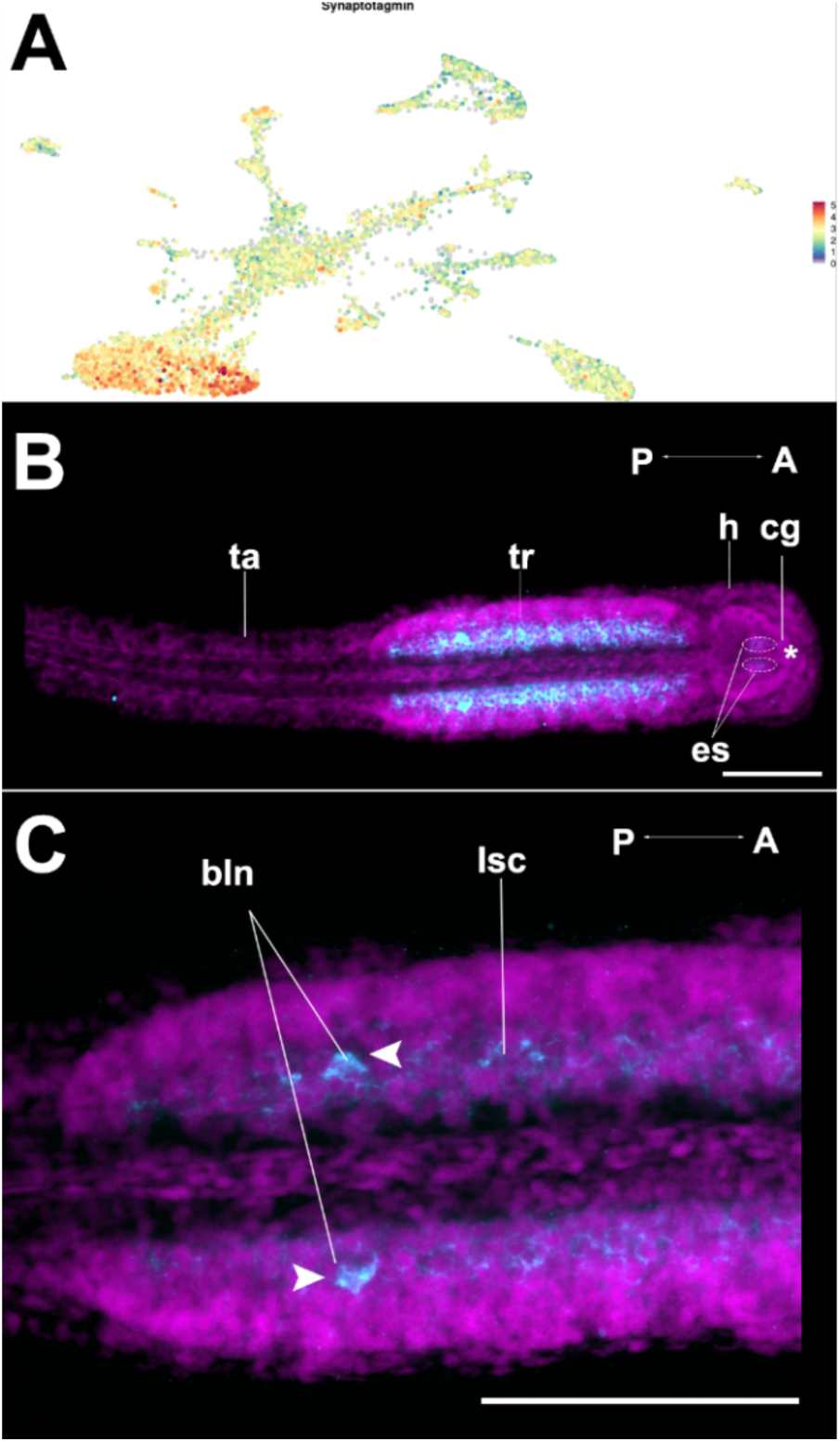
Cluster neurons 1 are identified by *synaptotagmin* in the *S. cephaloptera* hatchling (1dph). Gene expression is visualized with Fast-Blue (cyan), and nuclei with DAPI (magenta). **A.** Distribution of expression levels of the transcript encoding for *Sce-synaptotagmin* in the cell landscape recovered from the hatchling stage. **B.** Maximum projection of the ventral region of the hatchling, with a stronger expression of *Sce-synaptotagmin* in the trunk (tr) region. The expression domains in the head (h), correspond to the forming esophageal ganglia (circle dotted lines). **C.** Detail of the trunk region, showing the expression of *Sce*-*synaptotagmin* in the lateral somata cell clusters (lsc) in the ventral region of the trunk (arrowheads). Two additional expression domains match the location of the bilateral neurons (bln). The asterisk marks the location of the mouth opening. In the UMAP, red coloring indicates higher marker expression. Scale bars 100 µm. Additional abbreviations: ta, tail; cg, cerebral ganglion.

After hatching, the VNC of *S. cephaloptera* encompasses a variety of neuron types with distinguishable histological features in adults [8] [10]. Among them, cluster neurons 3 express the excitatory amino acid transporter *GLT-1* (Fig. 3A). ISH localizes these cells to the medial region of the lateral somata clusters, adjacent to the ventral neuropil (Fig. 3B-D). The expression pattern resembles two dotted strips that range the trunk across the longitudinal axis (Fig. 3B). Furthermore, Sce-*GLT-1^+^* cells were also found around the developing connectors between the cephalic ganglia and the ventral nerve center (Fig. 3D).

**Figure 3.**
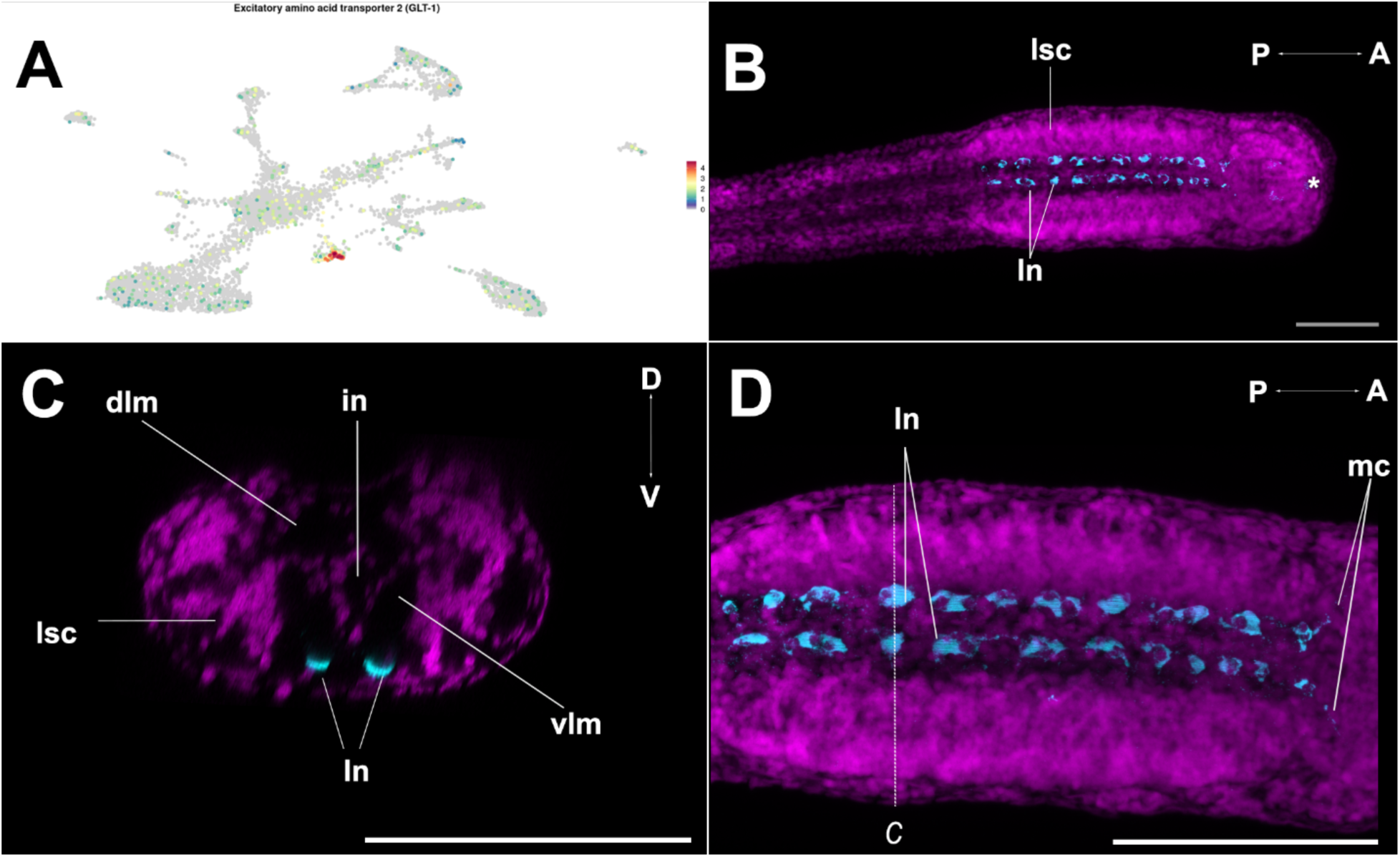
Expression of *excitatory amino acid transporter 2* (*GLT-1*) defining cluster neurons 3 in the hatchling (1dph) of *S. cephaloptera*. Gene expression is visualized with Fast-Blue (cyan), and nuclei with DAPI (magenta). **A.** Expression distribution of the transcript encoding for *GLT-1* across the cellular landscape of *S. cephaloptera*. **B.** Maximum intensity projection of the expression pattern obtained by ISH of *Sce-GLT-1*. The expression forms two parallel strips in the ventral region of the trunk, in the inner limit of the lateral somata cell cluster (lsc), within the cell bodies of large neurons (ln). **C.** The transverse section of the trunk region (referenced in D) shows two expression domains located in the ventral region of the lsc. **D.** Close-up of the trunk, displaying the expression domains in the anterior part of the trunk in the main connectives. The asterisk marks the location of the mouth opening. In the UMAP, red coloring indicates higher marker expression. Scale bars 100 µm. Additional abbreviations: in, intestine; dlm, dorsal longitudinal muscle; vlm, ventral longitudinal muscle.

The last neuronal cluster, neurons 4, is defined by neurons expressing a variety of receptors, including an *FMRFamide receptor,* together with one *GABA transporter* (GAT-2). Interestingly, this is the only neuronal cluster for which the transcription factor *Hox1* was identified as a gene marker, suggesting a possible Hox-mediated patterning role. Nevertheless, the location of those cells remains to be confirmed.

#### Muscle cells

Although 1dph hatchlings do not yet display predatory behavior, they possess a well-developed musculature primarily used for locomotion in response to external stimuli and escape from disturbances [10]. Two muscle cell-related clusters corresponding to distinct metacell populations were identified. The cell cluster muscle 1, featured canonical muscle markers such as *myosin heavy chain*, *regulatory myosin light chain* and *troponin I-4* (*TnI4*) [14][50], with *TnI4* expressed in approximately ∼60% of the cells of this cluster (Fig. 4A). The spatial distribution of the expression of *Sce-TnI4* encompasses the trunk and tail longitudinal muscles, as well as the vestibular muscles in the head region (Fig. 4B). Later in development, these cephalic muscles will be associated with forming a lateral plate to control the opening of the mouth. The cell cluster muscle 2 also expresses structural muscle markers such as *D-titin* and *twitchin.* However, compared to muscle 1, it exhibits a distinct differential expression of specific gene markers, such as *myosin-7*. Furthermore, its transcriptomic profile, as revealed by the Metacell analysis, suggests that muscle 2 represents a distinct muscle population. (Fig. 1D, E, F).

**Figure 4.**
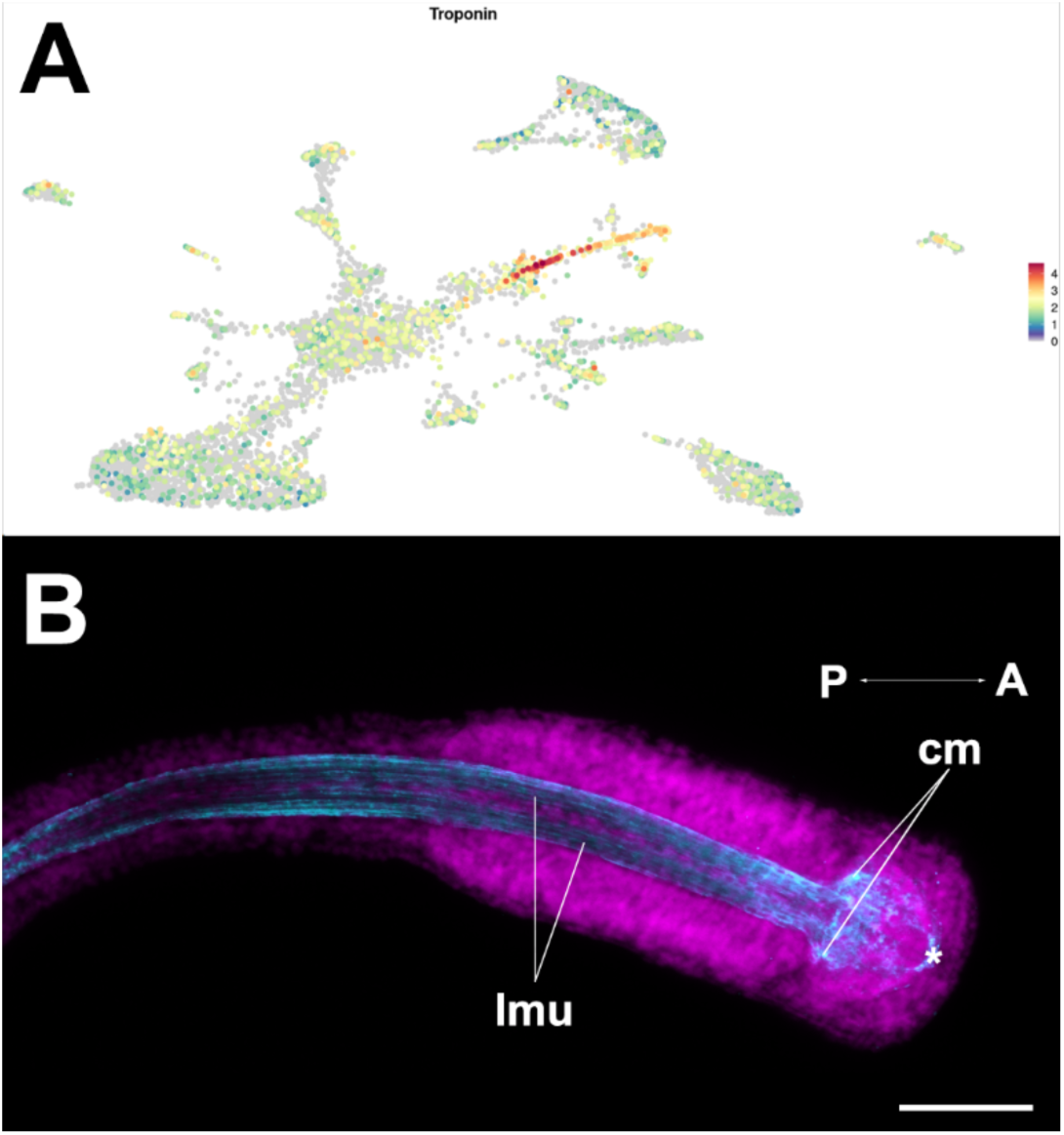
Cluster muscle 1 visualized by the expression of *troponin-1* in the hatchling (1dph) of *S. cephaloptera*. Gene expression is visualized with Fast-Blue (cyan) and nuclei with DAPI (magenta). **A.** Distribution of the gene expression of troponin-1 across the transcriptomic landscape of the cells recovered from the hatchling stage. **B.** Overview of *Sce-troponin-1* expression pattern in the hatchling, showing localization in the longitudinal muscles in the trunk and tail (lmu), and in the cephalic muscles (cm) of the vestibular region. The asterisk marks the location of the mouth opening. In the UMAP, red coloring indicates higher marker expression. Scale bars 100 µm.

#### Epidermal and gland cells

In chaetognaths, the epidermis exhibits a distinctive multilayered structure across multiple body regions [8]. Furthermore, the epidermis is also innervated and, from the hatchling stage onward, supports additional cell types located in the most distal layer of the ventral epidermis, as is the case for the adhesive papillae [8][10]. Although cells of the adhesive glands and the epidermis clusters share transcriptomic similarities (Fig. 1D), they remain distinct cell types, distinguishable by differential transcript-expression patterns.

Epidermal markers such as *Sce-fibrillin*-*1* are expressed in the distal epidermal cells across the entire specimen (head, trunk, and tail) (Fig 5A-D). Strong expression was observed in regions where adhesive papillae are abundant, for instance, in the anterior-most part of the ventral side of the head (Fig. 5B), and the transition zone between trunk and tail (Fig. 5C). *Sce-calmodulin*, another epidermal marker, displays a similar expression pattern as *Sce-fibrillin-1*. It is expressed in the distal-most layer of the epidermis (Fig. 5E-G) and exhibits a stronger signal in the cephalic adhesive papillae and posterior trunk (Fig. 5F, G).

**Figure 5.**
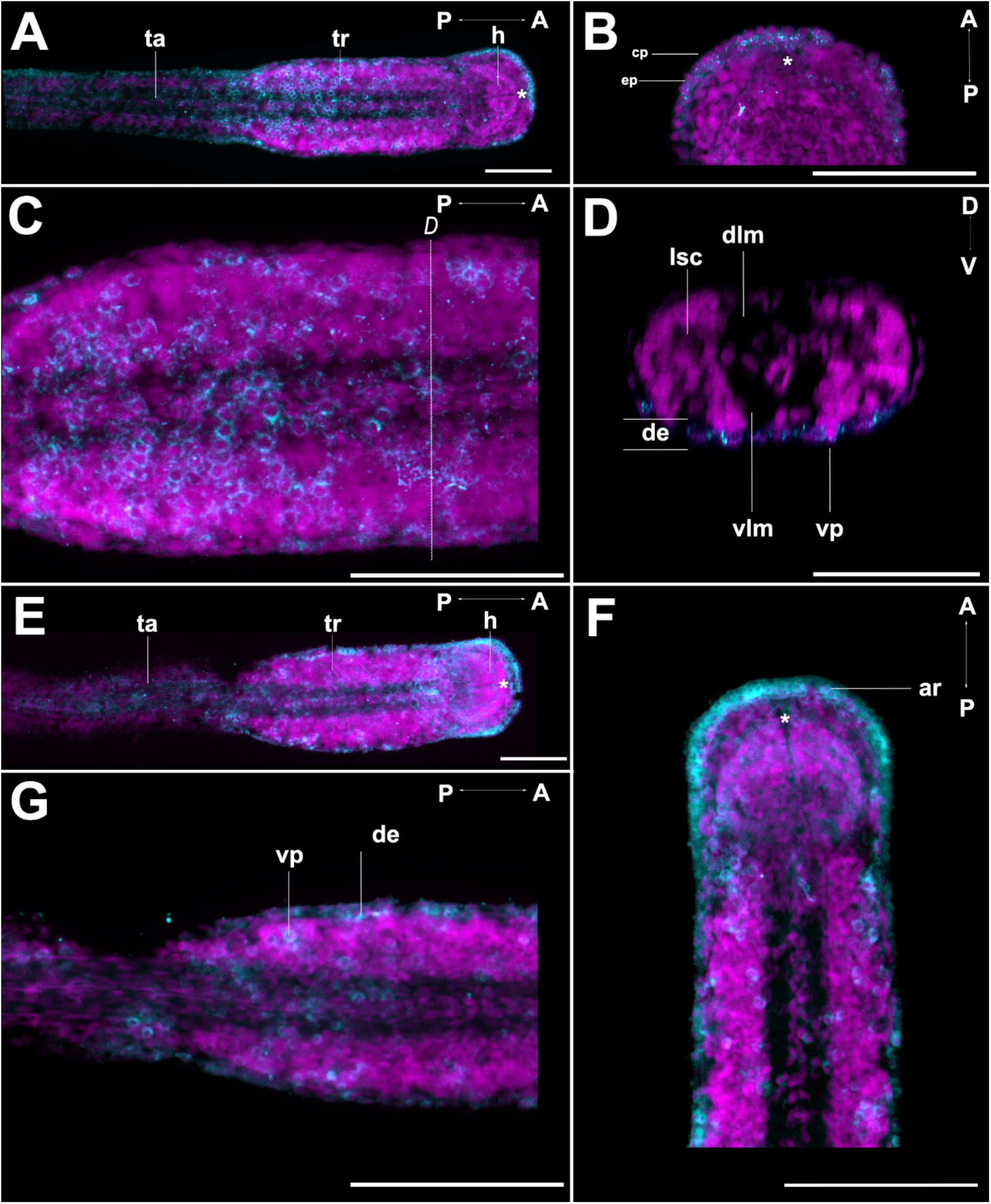
Expression of the epidermal cell cluster maker genes, *fribrillin-1* (A-D) and *calmodulin* (E-G) in hatchlings 1dph of *S. cephaloptera*. Gene expression is visualized with Fast-Blue (cyan) and nuclei with DAPI (magenta). **A.** Overview of the expression pattern that involves the entire surface of the hatchling: head (h), trunk (tr), tail (ta). **B.** Close-up of the head region where the expression of *Sce*-*fibrillin-1* is stronger in the anterior rim of the head, involving distal epidermal cells (ep) and cephalic papillae (cp). **C.** Close-up of the trunk where expression is stronger in the posterior region **D.** Transverse section (referenced in C), depicting that the expression of *fibrillin-1* involves the distal epidermis (de) and the ventral papillae (vp). **E.** Overview of the expression of *Sce-calmodulin* in the hatchling, which resembles the expression of *Sce*-*fibrillin-1*. **F.** Close up in the head and the anterior part of the trunk, where the expression of *Sce-calmodulin* includes the anterior rim (ar). **G.** Detail of the anterior part of the tail where the expression is related to both distal epidermal cells and ventral papillae. The asterisk marks the location of the mouth opening. Scale bars 100 µm. Additional abbreviations: dlm, dorsal longitudinal muscle; lsc, lateral somata cluster; vlm, ventral longitudinal muscle.

Cells with increased expression of *fibropellin-1* group together with those cells of the proposed adhesive gland cluster (Fig. 6A). In 1dph old individuals, *Sce-fibropellin-1*-expression is confined to individual pyramidal-shape cells localized in the most distal layer of epidermal tissue, distributed in the ventral side, near the septum between trunk and tail, in the trunk, and the cephalic ring in the head (Fig. 6B, C, D). To further confirm the identity of this cell type, the expression pattern of *Sce*-*fibropellin-1* was examined in 5 days after hatching individuals (5dph) (Fig. 6E, F, G) and in adults (Fig. 6H). In 5dph specimens, the expression persists in the ventral region of the animal but shifts towards the posterior end of the tail (Fig. 6E). Additionally, the expression of *Sce*-*fibropellin-1* in the anterior region of the head is almost absent, with a few cells of the anterior rim showing slight expression (present at 1dph) (Fig 6G). Adults express *Sce-fibropellin-1* in the mid-anterior part of the tail, where aggregates of cells, instead of individual cells, now express this gene marker (Fig 6H).

**Figure 6.**
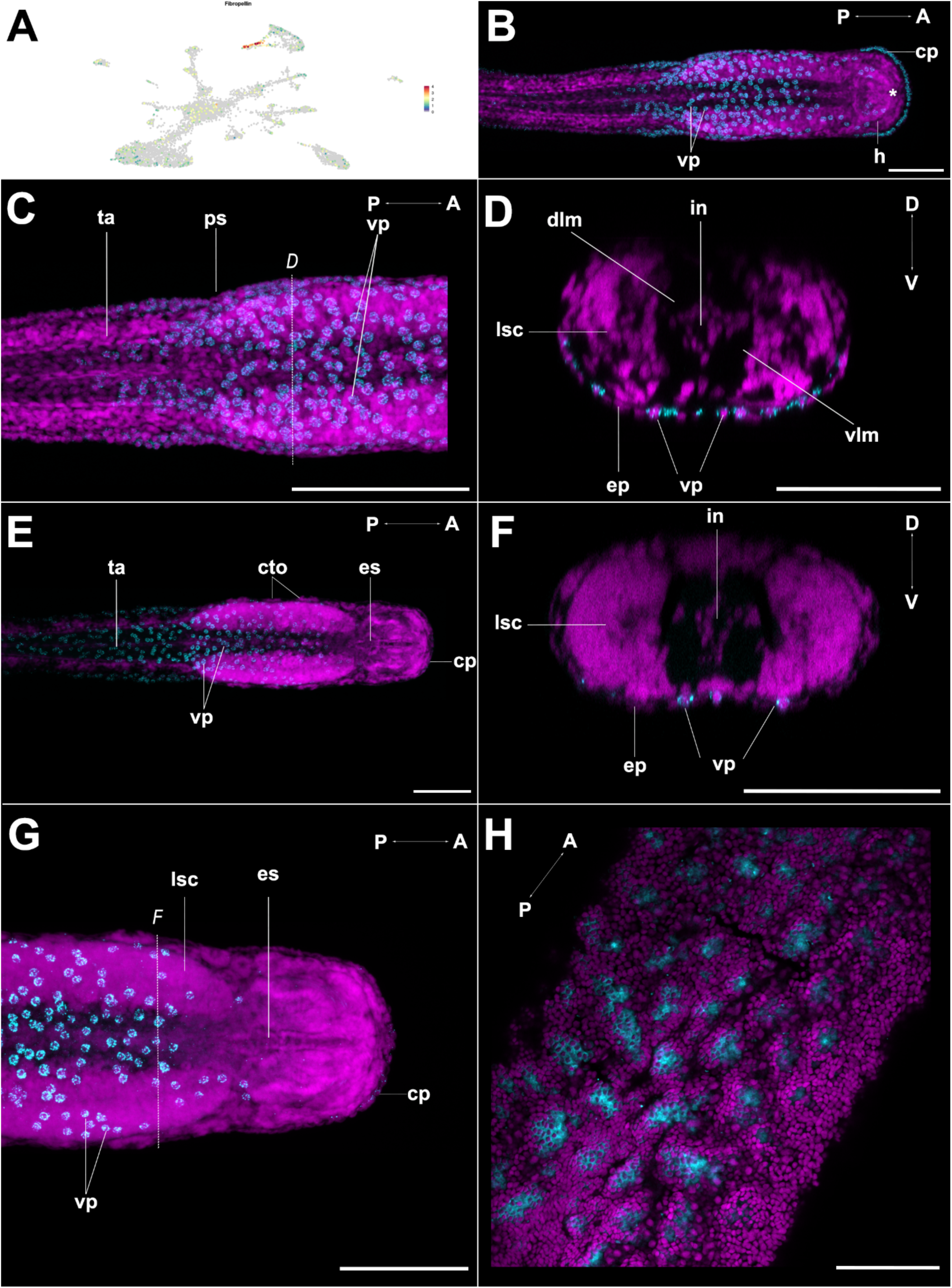
Expression of fibropellin *–1* highlights the adhesive gland cell cluster in hatchlings 1dph (B-D), 5dph (E-G), and adult (H) of *S. cephaloptera*. Gene expression is visualized with Fast-Blue (cyan) and nuclei with DAPI (magenta). **A.** Distribution of cells expressing higher levels of *Sce-fibropellin-1* within the hatchling cell-transcriptomic landscape. **B.** Overview *Sce*-*fibropellin-1* expression pattern in 1dph, localized to the anterior cephalic adhesive papillae (cp), in the ventral papillae (vp) of the trunk, and the most anterior part of the tail. **C**. Close-up in the ventral region of the transition between trunk and tail, where cells on both sides of the posterior septum (ps) exhibit expression of *Sce-fibropellin-1*. **D.** Transverse section (indicated in C), revealing that the *Sce-fibropellin-1* expression is confined to discrete ventral papillae cells in the distal epidermis, while normal epidermal cells (ep) lack transcript signal **E.** Overview of *Sce-fibropellin-1 expression* in a 5dph specimen of *S. cephaloptera*, showing a posterior shift of expression in the tail (ta). **F.** The transverse section (referenced in G) displays expression in individual adhesive ventral papillae cells. **G.** Close up to the anterior part of the trunk and head of the 5dph, showing the near absence of expression in the anterior rim. **H.** Overview of *Sce-fibropellin-1* expression in the anterior part of the tail of the adult stage, where the transcript signal now corresponds to clusters of multiple cells. In the UMAP, red coloring indicates higher marker expression. The asterisk marks the location of the mouth opening. Scale bars 100 µm. Additional abbreviations: cto, ciliary tuft organ; dlm, dorsal longitudinal muscle; es, esophagus; h, head; in, intestine; lsc, lateral somata cluster; ta, tail; vlm, ventral longitudinal muscle.

In addition, the adhesive cell gland cluster markers include genes encoding for proteins that may directly mediate the attachment mechanism in the hatchling stage. For instance, approximately 95% of the cells in this cluster express a putative *extracellular protease inhibitor*, along with *Snaclec coagulation factor IX*, *calmodulin*-like proteins, *zinc-metalloprotease*, and *laminin-subunit alpha-2*.

#### Ciliary receptors

In *S. cephaloptera,* the epidermis contains various ciliary sensory organs [8][51]. Metacell and Seurat analyses identified three distinct cell populations enriched in cilia-related genes, such as *dynein* [14]. These ciliary receptor populations differ both from the epidermis cell cluster and each other in their transcriptomic profiles (Fig. 1D). Cells of the cluster ciliary receptors 1 (Fig. 7A) express *dynein beta chain* and form the ciliary tuft organs, which are located longitudinally in both dorsal and ventral regions (Fig. 7B). Moreover, the expression domain of this marker includes the receptors present along the lateral margins of the head, trunk, and tail (Fig 7B, C). Interestingly, *Sce-dynein beta chain* expression is absent in the ciliary receptors located closer to the longitudinal midline. Furthermore, the expression of this gene is predominantly clear in the sensory cells of each ciliary tuft organ, which are embedded into the multilayered epidermis (Fig. 7D).

**Figure 7.**
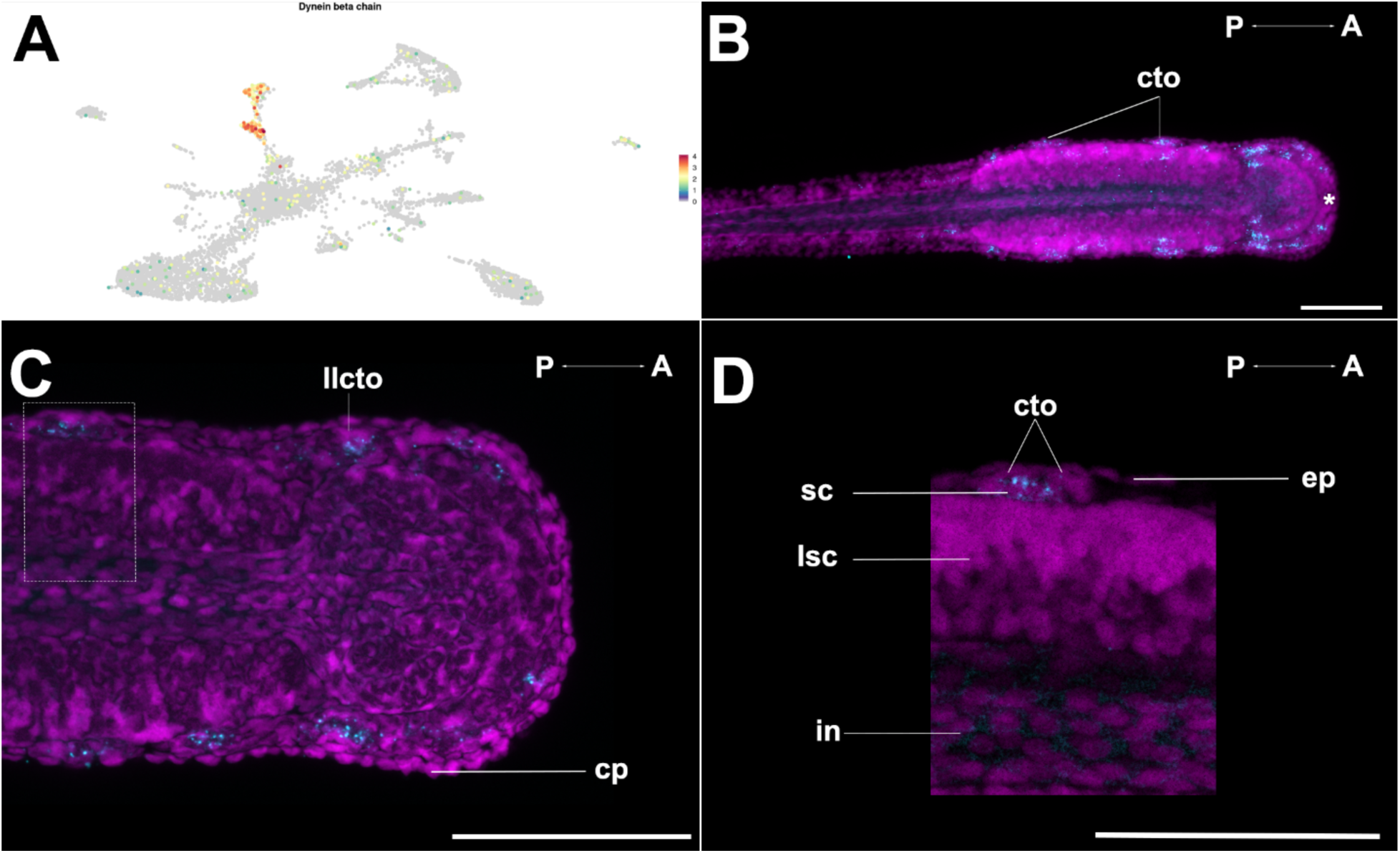
Distribution of the transcripts of *dynein beta chain* reveals cluster ciliary receptors 1 in the hatchling of *S. cephaloptera*. Gene expression is visualized with Fast-Blue (cyan) and nuclei with DAPI (magenta). **A.** Distribution of the cells expressing *Sce-dynein beta chain* in the transcriptomic landscape of the hatchling stage. **B.** Maximum projection overview of the expression pattern of *Sce*-*dynein beta chain,* showing cell clusters that distribute longitudinally in the distal part of the epidermis (dorsally and ventrally), corresponding to ciliary tuft organs (cto). **C.** Close-up in the anterior part of the hatchling, where the gene expression is visible in the lateral line ciliary tuft organs (llcto). **D.** Detail of the expression of *Sce*-*dynein beta chain* within one ciliary tuft organ (dotted rectangle in C), displaying expression in the sensory cells (sc), which are distally positioned relative to the lateral somata cluster (lsc). The asterisk marks the location of the mouth opening. In the UMAP, red coloring indicates higher marker expression. Scale bars 100 µm. Additional abbreviations: ep, epidermis; in, intestine.

Ciliary receptor cluster 2 differs markedly from cluster 1, showing only limited transcriptomic similarity and thereby constituting a distinct cell population (Fig. 1D, E, F). Cells of cell cluster ciliary receptors 2 express genes related to mechanosensory processes such as *hemicentin-1* [52], and microtubule-dependent transporters essential for the formation and maintenance of cilia such as *kinesin*-like *protein* [53]. Notably, this cell population is the only ciliary group of metacells expressing *polycistin-1*, a gene related to mechanosensory cells. Considering the above, this cell type could be potentially associated with additional ciliary receptors present in the epidermis of the adult.

The corona ciliata are two concentric rings of ciliated sensory cells that is located on the dorsal side in the trunk-head transition region. Constituting cells for this organ form a distinct cell cluster that expresses a variety of genes associated with ciliary cells and sensory cells, such as *dynein heavy chain-like protein 2* [14], and *roundabout homolog 2* [54] (*robo2*), respectively (Fig. 8A). In *S. cephaloptera* hatchlings, *robo2* expression is concentrated in the corona ciliata region (Fig. 8B, C), with lower expression levels in the developing main connectives (Fig. 8D).

**Figure 8.**
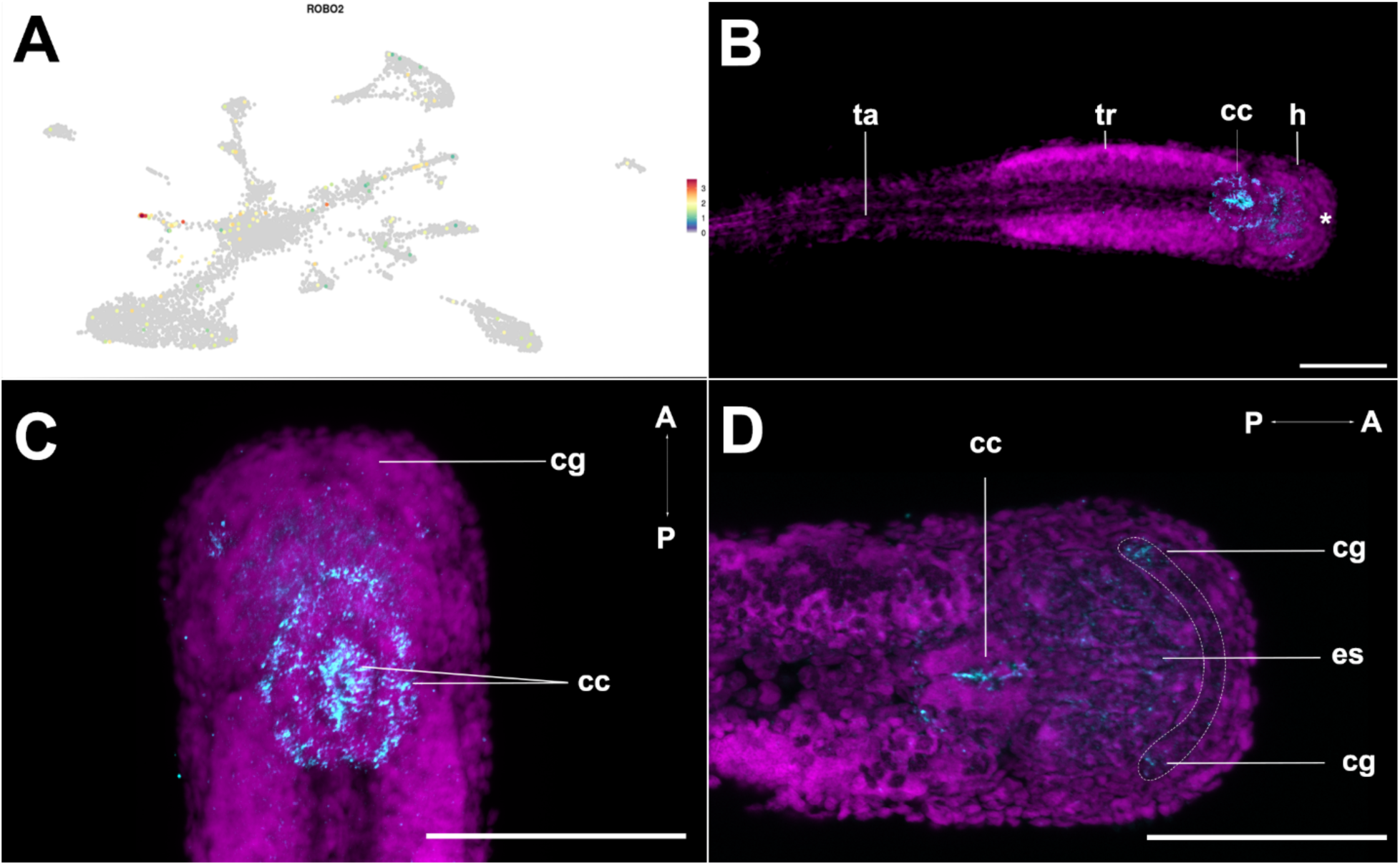
Cells belonging the cell cluster corona ciliata are distinguished by the expression domains of *robo2* in the hatchling of *S. cephaloptera*. Gene expression is visualized with Fast-Blue (cyan) and nuclei with DAPI (magenta). **A.** Pattern of *Sce*-*robo2* expression across the transcriptomic landscape of cells analyzed at the hatchling stage. **B.** Overview of *Sce*-*robo2* expression pattern, showing signal dorsally, in the transition between trunk (tr) and head (h), corresponding to the location of the corona ciliata (cc). **C.** Maximum projection of the expression pattern of *Sce-robo2* in the anterior region, in this view the two concentric rings of the corona ciliata are visible. **D.** Close up of the anterior part of the head region, where expression is bilaterally observed in the distal part of the forming cerebral ganglion (cg) (dotted lines), being stronger in the presumed developing coronal nerves. The asterisk marks the location of the mouth opening. In the UMAP, red coloring indicates higher marker expression. Scale bars (A-C) 100 µm D 50 µM. Additional abbreviations: es, esophagus; ta, tail.

#### Sperm duct

Another cell cluster identified for the hatchling of *S. cephaloptera* possesses cells that contribute to the development of the male reproductive system. Chaetognaths are protandric hermaphrodites, and the male reproductive organs develop first, followed by the female organs. In adults, the tail coelom contains the spermatogonial masses and connects the testis with the sperm vesicles through sperm ducts, endowed with ciliated cells [8]. Marker genes for this sperm duct cell cluster are linked to ciliated-cell activity, as is the case of *rootletin* and *kinesin*-like protein (*KIF21A*) [55]. The expression pattern of one marker with no described ortholog (*Sce-45089*), includes 67% of the cells of this cluster (Fig. 9A). Its expression pattern by ISH is restricted to the posterior-most tail region, close to the forming caudal fin (Fig. 9B, C). In addition, the signal extends anteriorly in two adjacent strips (Fig. 9C, D), up to the anterior part of the tail, near the posterior septum, presumably in the region of the male germ cells (data not shown).

**Figure 9.**
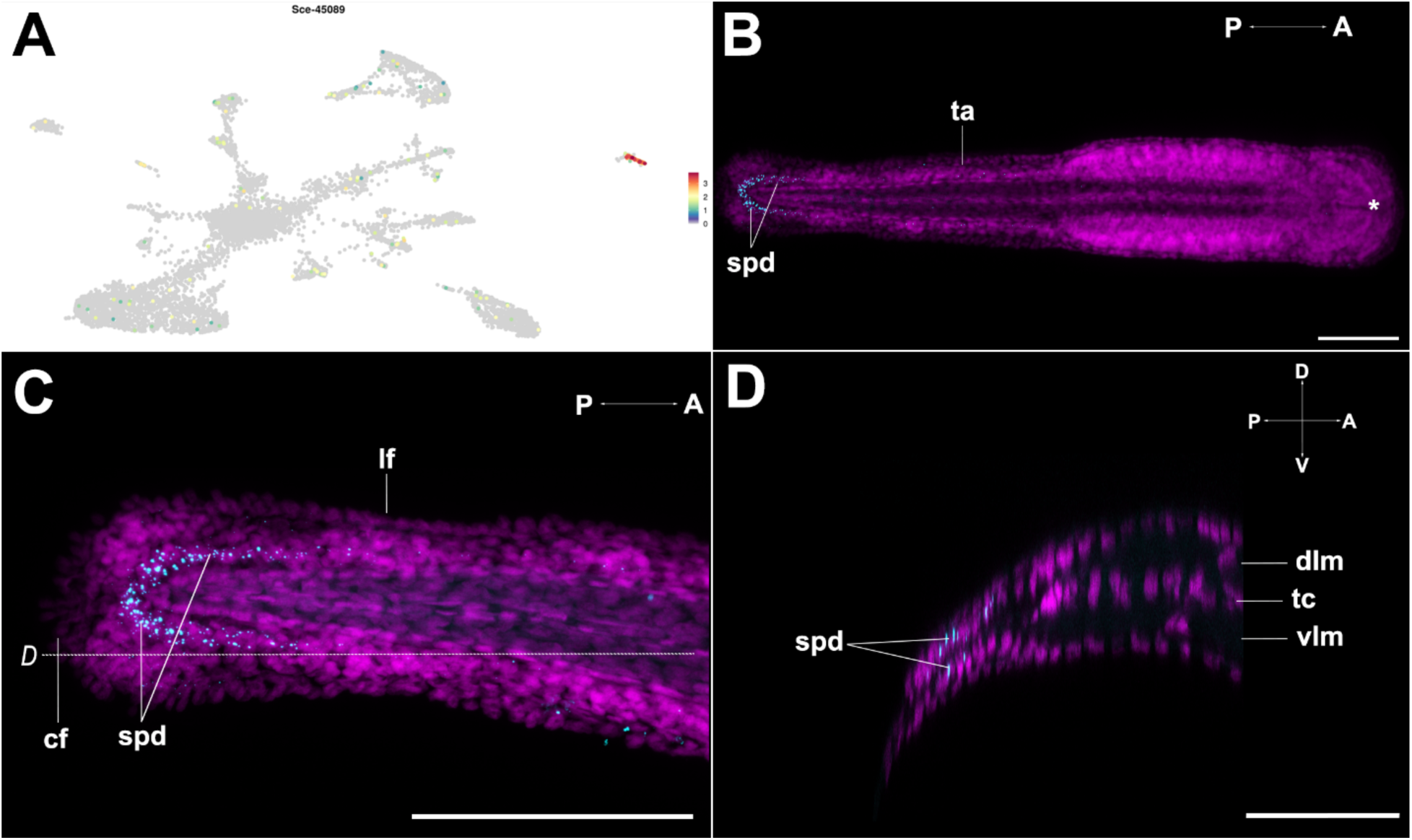
Expression of *Sce-45089* (spermduct-associated) marker in the hatchling of *S. cephaloptera*. Gene expression is visualized with Fast-Blue (cyan) and nuclei with DAPI (magenta). **A.** Expression of *Sce-45089* visualized in the transcriptomic profile of cells from the hatchling stage. **B.** Overview of the expression pattern of *Sce-45089*, present in the most posterior part of the tail involving the forming sperm duct (spd) **C.** Close-up of the posterior region of the tail, showing the transcript’s signal in the sperm duct near the caudal fin (cf). **D.** Longitudinal view (referenced in C), where the two adjacent expression domains in the sperm duct are visible. The asterisk marks the location of the mouth opening. In the UMAP, red coloring indicates higher marker expression. Scale bars 100 µm. Additional abbreviations: dlml, dorsal longitudinal muscle; f, lateral fin; tc, trunk coelom; vlm, ventral longitudinal muscle.

#### Germ cells

Germ cells are clearly identifiable in *S. cephaloptera*, and their transcriptomic profile includes genes involved in germ cell formation of other organisms, such as *APOBEC1 complementation factor* (Fig. 10A), and genes involved in lipid metabolism, such as *apolipophorin-2* [56][57]. This cell type encompasses a few cells, and Sce-*APOBEC1* expression is located in the transition zone between trunk and tail (Fig. 10B, C). The ventral expression domains in the anterior-most part of the trunk correspond to the female germ cells, and the signal coming from the ventroanterior part of the tail is related to the forming male germ cells (Fig. 10C).

**Figure 10.**
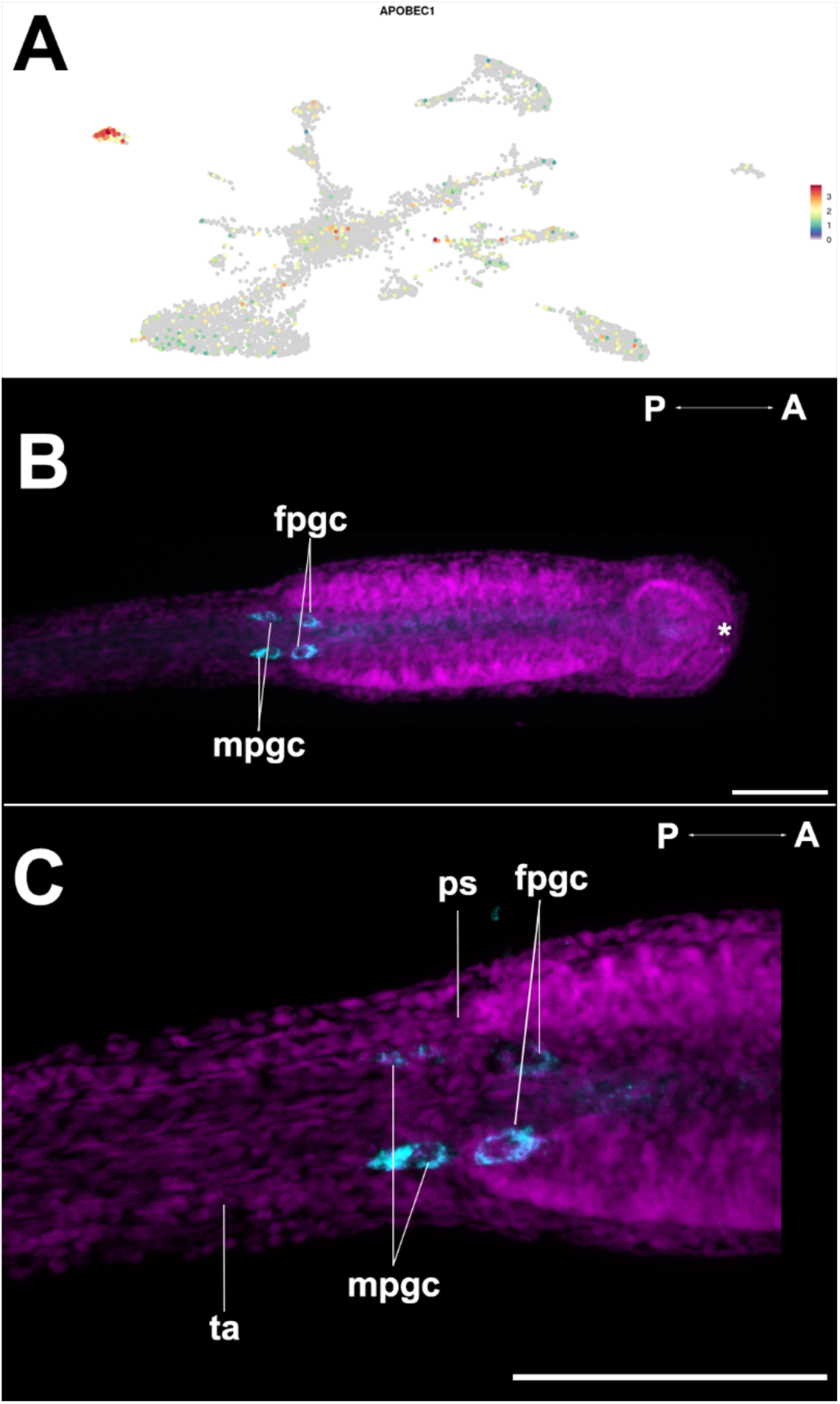
Identification of germ cells by the expression pattern of *APOBEC1* in the hatchling of *S. cephaloptera*. Gene expression is visualized with Fast-Blue (cyan) and nuclei with DAPI (magenta). **A.** Distribution of the expression of *Sce*-*APOBEC1* transcript in the cells recovered from the hatchling stage. **B.** Overview of *Sce*-*APOBEC1* expression pattern, with signal localized ventrally in the transition between trunk and tail, corresponding to the male germ (mpgc) and female germ cells (fpgc). **C.** Detail in the posterior septum (ps), displaying the distribution of the germ cells in the hatchling stage. The asterisk marks the location of the mouth opening. In the UMAP, red coloring indicates higher marker expression. Scale bars 100 µm. Additional abbreviations: ta, tail.

#### ECM-related cells

In *S. cephaloptera,* the most internal/ proximal cells of the multilayered epidermis are organized on top of the ECM. Stacked fibrillous content is enriched in this region, conforming to the basal laminae [8]. Two different cell types constitute the basal laminae of chaetognaths. While the outer area is made up of epidermal cells, the inner region is built from mesoderm derivatives [8]. In this context, by examining the gene markers for the putative cluster of ECM-related cells, it was found that this cell population has high expression levels of *collagen alpha-1* and *nidogen*, which are known components of the basal laminae [58]. Notably, approximately 97% of these cells express the gene encoding for *serrate* (the homolog of *jagged-2*), known to be involved in the *notch* signaling pathway, which may be modulating a *serrate-notch*-dependent cell fate determination [59].

#### Immune cells

Marine invertebrates rely on innate immune responses to defend against infections. In this context, an analysis of the putative immune cell cluster reveals a high abundance of various receptors capable of recognizing external pathogens, such as *scavenger receptors* and *c-type lectin* domains. Additionally, a gene encoding for the transmembrane family member 1 protein (*SIDT1*), which is involved in the transport of interference RNA (RNAi) [60], was identified in approximately 85% of the cells within this cluster. These components may facilitate the targeted elimination of the genetic material of invading agents, such as viruses.

## Discussion

### Neuronal diversity

At hatching, the nervous system of *S. cephaloptera* already exhibits many of the ganglionic anlagen seen in adults, with developing ganglia in both the cephalic region and the nascent ventral nerve center in the trunk [10][13]. The developing cerebral ganglion in the head region connects to the ventral nerve center in the trunk. Furthermore, two other cephalic ganglia (vestibular and esophageal) are formed by cells with a distinct transcriptomic profile [13]. Considering the above, the different neuronal cell populations identified in the hatchling stage are likely to be either progenitors of later-stage structures or fulfill crucial functions that allow the animal to respond to stimuli within the first few days post-hatching [10].

Among the four neuronal clusters identified, neurons 1 and neurons 2 share extensive transcriptomic similarity, for instance, both of them exhibited in over 90% of their cells expression of *glycine receptor subunit alpha-3* (GlyaRa3), yet they differ in key gene markers. Neurons 1 is defined by the expression of genes related to synaptic activity, such as a homolog of *alpha 1A adrenergic receptor* [61] and *glutamate receptor U1* (*Kainate*) [62]. On top of that, those cells contain in their transcriptomic program, *forkhead box P1* (*FoxP1*), which has been linked to neuronal development in the early stages of *Drosophila melanogaster* [63], and it has been identified in the pallial neurons associated with glutamatergic activity in the lamprey *Petromyzon marinus* [64] and in the annelid *Capitella teleta* [65]. Furthermore, in *Ciona intestinalis*, glutamatergic neurons are related to different types of sensory neurons, including photoreceptors, gravity-sensitive antennae, and epidermal cells [66]. The co-expression of *GlyRa3* and *Kainate* suggests that neurons 1 may integrate glycinergic and glutamatergic input as observed in neurons containing NMDA receptors in *D. melanoganster* [67]. However, the expression of homologs of adrenergic receptors that, in invertebrates, typically associated with octopaminergic signaling [68], suggests that neurons 1 may represent a cell population integrating multiple signaling pathways that would require further validation to clarify the interplay of these neurotransmitters.

In the case of cluster neurons 2, the cells express transcripts such as *neuronal acetylcholine receptor subunit alpha-6* and *short transient receptor potential channel-6*. In chaetognaths, acetylcholine acts as the primary neuromuscular transmitter [8]. The axons do not directly contact the muscles; instead, a thick extracellular matrix (ECM) separates the presynaptic terminals from the locomotive muscle fibers. The axons terminate at the epidermal side of the ECM, and from there, acetylcholine diffuses across to the muscle cells.[8]. Furthermore, the expression of cholinergic markers (*ChAT* and *VAChT)* has already been described to exhibit expression domains in the “inner” cells of the lateral somata clusters of the VNC in hatchlings of *S. cephaloptera* [13]. Moreover, a similar cell type in the sea urchin *Strongylocentrotus purpuratus* has been described as a cholinergic cluster [69]. Collectively, these results match the expression pattern observed from the ISH of the marker *Sce*-*synaptotagmin* (Fig. 2B, C) and support the idea that neurons 2 correspond to a cholinergic population. Nevertheless, it is still to be elucidated if these cells have expression of the transcript combination *pax6*, *nkx6*, and *hb9* that would allow identifying them as potential cholinergic motor neurons [69].

The cluster neurons 3 has associated markers such as the *glycine receptor subunit-2* and *glycine receptor subunit-3* (*GlyR*). This receptor type has been reported to modulate pattern generation movement that allows coordinated swimming in the ascidian *C. intestinalis* [70]. In this regard, *Ci-GlyR* exhibits an expression pattern along the tail and the motor ganglion in the hatched larva stage [70]. On the other hand, in the hatchling of *S. cephaloptera,* the expression domains of *Sce*-*GLT1,* a marker for neurons 3, are located ventrally in the trunk region (Fig. 3C). Interestingly, both expression patterns (*Ci-GlyR* and *Sce-GLT1*) resemble discontinuous lines located towards the anterior region of the animal (Fig. 3D). Furthermore, cluster neurons 3 has expression of non-glycine-related metabolism proteins, such as a glutamate receptor. For this reason, this cell population could not be considered strictly glycinergic neurons but rather regulated by their activity [71] [72]. On top of this, *acetylcholinesterase* is also a marker for this cell population, indicating that this cell type may be exposed to cholinergic input [73]. These findings suggest that cells in this cluster are neurons that drive a regulatory role between cholinergic and glycinergic neuron pathways [74].

The last neuronal cell type identified corresponding to neurons 4 exhibits a *sodium and chloride-dependent GABA transporter 2* (*GAT-2*), which has been reported to participate in the uptake and recycling of the neurotransmitter GABA in *C. elegans* at GABAergic synapses [75]. In addition, neurons 4 exhibits receptor homologs for the neuropeptide FMRFamide. In this sense, previous research documents the distribution of RFamide-related peptides in hatchlings of *S. cephaloptera* in the VNC, involving lateral cells, dorsal cells, and longitudinal fiber bundles in the trunk [10]. Moreover, this cell cluster shows expression of the transcription factor *Hox1*, which could potentially regulate the location of cells within the VNC. In this regard, the distinctive location of a median Hox gene (*Med4*) in the VNC *of S. cephaloptera* has been already observed [76]. Considering the above, it is likely that this cell population is defined by neurons modulating motor behaviors [10].

### Epidermal and gland cells

The epidermis of *S. cephaloptera* supports various cell types, including ciliary receptors and adhesive glands (Figs. 6-8) [8]. In this regard, the epidermal cell cluster shows high expression of structural markers that may provide support to such structures as *collagen alpha-1*(*IV*), *collagen alpha XXII*, and *intermediate filament protein-B*. Notably, collagen distribution in the integument of marine invertebrates has been described as a common feature among echinoderms, as is the case of *Stichopus herrmanni,* which provides structural support for additional systems such as papillae [77] [78]. In this context, collagen fibers allow the echinoderms to have resistant connective tissue under tensile stress [79]. This stability would be particularly relevant for an organism such as *S. cephaloptera,* which exhibits temporary attachment and requires moderate resistance against disturbances.

The multilayered epidermis of *S. cephaloptera* also houses a neuronal plexus projected from the VNC in the trunk [10]. In this sense, the epidermis cell cluster was observed to express a different isoform of the synaptic marker *synaptotagmin*. Given the foregoing, it has been observed that the regions of the epidermis below the adhesive glands exhibit more abundance of neuronal processes. This observation has led to hypothesize a nervous system modulation of the attachment in *S. cephaloptera* [8], a phenomenon also observed in the tunicate *C. intestinalis* [80]. On the other hand, the close (spatial) relationship between the epidermis and the adhesive glands in *S. cephaloptera* observed histologically in the adult [8] may explain the high degree of similarity observed in their transcriptomic profiles (Fig. 1D). This similarity of their transcriptomes also accounts for the higher intensity of the expression patterns from *Sce-fribillin1* and *Sce-calmodulin* in the regions enriched in adhesive cells (Fig. 6).

After hatching, individuals of *S. cephaloptera* primarily attach through the anterior part of the body using the adhesive papillae [10]. In adults, these cells have elongated shape and swollen tips enriched in secretion granules, which may contain some of the molecular agents involved in the attachment process [8]. In addition, in the adult, those cells form clusters of around six cells and localize mainly in the distal ventral epidermis close to the septum between the trunk and tail [9]. Nevertheless, until now, the molecular agents that may mediate the attachment process in *S. cephaloptera* have been unexplored. The present study observed that this cell type is defined by the expression of two protease inhibitor-like proteins, which are expressed in more than 90% of the cells in this cell population. In this regard, these kinds of proteins have been described to mediate attachment in the pedal disc of the cnidarian *Exaiptasia pallida* [81] and to be present in the sticky silk of the larval stage of the caddisfly *Bombyx mori* [5]. On the other hand, both putative protease inhibitors found in the cell cluster contain a signal peptide in their sequence, which leads to the hypothesis that *S. cephaloptera* relies on a secretion with protein content to adhere to different substrates. These findings provide further evidence for the convergent evolution hypothesis for marine invertebrate adhesives, as distantly related marine invertebrates appear to have independently arrived at similar biochemical solutions [2].

Additional markers for the adhesive gland cell type, such as coagulation factor IX, provide further similarities to the adhesive systems reported in *Exaiptasia pallida* and *Paracentrotus lividu*s, both of which express coagulation factor-like proteins in their secreted glues [81] [82]. Moreover, the adhesive cell cluster of *S. cephaloptera* also expresses calcium-binding domain-containing proteins, which have been described as part of the non-permanent adhesion mechanism of *Patella vulgata* [83]. In addition, this cell population in *S. cephaloptera* expresses *synaptotagmin 7*, which provides further support to the idea of a ventral nerve center regulation of the attachment process. Furthermore, considering that there was no evidence of a unique neuron-cell type related to the attachment process, it is hypothesized that the adhesive gland cells mediate themselves the communication with the VNC, presumably without a dedicated “adhesion neuron” population.

From a developmental perspective, the expression pattern of *Sce-fibropellin-1* was observed during the early stages, defining individual adhesive cell glands (Fig. 6B-D), and later in adults, it was visualized in clusters of cells (Fig. 6H). In this regard, it may be that the clusters of cells described by Müller et al., 2019 [8], occur only in late development, as they may be necessary to mediate the attachment of a larger specimen. Furthermore, the gene expression pattern of *Sce*-*fibropellin+* cells matches the behavior of *S. cephaloptera* described by John in 1933 [9]. In this context, he described that 24 hours after hatching the individuals do not feed and tend to group close to the hatching site, which explains the expression of the attachment gene marker in the mouth region (Fig. 6B). However, after 5 days of development the hatchlings of *S. cephaloptera* are capable of feeding (by lifting the head to capture their prey), and that is why the expression in the most anterior part of the head is abolished (Fig. 6G). Moreover, it was clear that in adults, the expression pattern in the tail region involves multiple cells (Fig. 6H), which matches the TEM description of Müller et al., 2019 [8]. In addition, it is relevant to consider the architecture of the *Sce-fibropellin-1* protein as it contains epidermal growth factor conserved domains (EGF), which is one of the hallmarks of the adhesive protein *Mlig-ap1* observed in the flatworm *Macrostomum lignano* [84]. Considering the above, here it is provided support for the similarities between adhesion mechanisms across taxa. Nevertheless, it is essential to keep in mind how proteins are built and that the presence of conserved domains in a diversity of clades speaks more of independent evolutionary processes shaped by similar environmental pressures, exemplifying convergent evolution in marine invertebrate non-permanent attachment systems [2].

### Conclusion

The use of single-nuclei transcriptomics enabled the characterization of the cellular complexity in hatchlings of *S. cephaloptera*, with a particular focus on its adhesive and nervous systems. Parallel transcriptomic and *in-situ* hybridization analyses shed light on the identification of adhesive gland cells distinguished by protease inhibitors and structural proteins observed in other marine invertebrate adhesives, underscoring the hypothetical convergent evolution of temporary attachment mechanisms. Together, these findings provide a broad molecular and cellular framework for understanding early developmental programs and organ system architecture of *S. cephaloptera*

## Author contributions

TW and CB conceived the idea and planned the project. TW secured funding for the project. CB and JFO collected the specimens for genome sequencing. JFO performed the DNA high-molecular weight extraction. CB performed RNA extraction from multiple stages for Iso-seq sequencing. CB and JM performed the genome assembly. CB and JM performed the transcriptome assembly and genome annotation. CB performed the single-nuclei analyses and validated cell types by *in situ* hybridization experiments. CB wrote the first version of the manuscript. TW contributed to all subsequent versions. All authors read, commented and finally approved the final version of the manuscript.

## Information on Funding

This research was supported, in whole or in part, by the Austrian Science Fund (FWF) [P34665]. CB and TW are grateful to the Vienna Doctoral School of Evolution and Ecology for funding the PhD position of CB. The authors thank the Faculty of Life Sciences of the University of Vienna (Austria) for financial support.

## Acknowledgements

Leslie Pan (EMBL Heidelberg, Germany) is acknowledged for valuable advice concerning the single-nuclei sequencing protocol. The authors also would like to thank Nikolaus Papadopoulos for his suggestions regarding genome assembly and single-nuclei analyses, the Faculty of Life Sciences at the University of Vienna for providing financial support, and the team from the Life Science Compute Cluster (LiSC) at the University of Vienna for their assistance with software usage.

## Notes

### Competing Interest Statement

The authors have declared no competing interest.

## References

1. Dodou, D., Breedveld, P., de Winter, J. C. F., Dankelman, J., & van Leeuwen, J. L. (2011). Mechanisms of temporary adhesion in benthic animals. Biological Reviews, 86(1), 15–32. 10.1111/j.1469-185X.2010.00132.x

2. Delroisse, J., Kang, V., Gouveneaux, A., Santos, R., Flammang, P. (2023). Convergent Evolution of Attachment Mechanisms in Aquatic Animals. In: Bels, V.L., Russell, A.P. (eds) Convergent Evolution. Fascinating Life Sciences. Springer, Cham. 10.1007/978-3-031-11441-0_16

3. Johnson, C. J., Razy-Krajka, F., Zeng, F., Piekarz, K. M., Biliya, S., Rothbächer, U., & Stolfi, A. (2024). Specification of distinct cell types in a sensory-adhesive organ important for metamorphosis in tunicate larvae. PLoS biology, 22(3), e3002555. 10.1371/journal.pbio.3002555

4. Liang C, Strickland J, Ye Z, Wu W, Hu B and Rittschof D (2019) Biochemistry of Barnacle Adhesion: An Updated Review. Front. Mar. Sci. 6:565. doi: 10.3389/fmars.2019.00565

5. Davey, P. A., Power, A. M., Santos, R., Bertemes, P., Ladurner, P., Palmowski, P., Clarke, J., Flammang, P., Lengerer, B., Hennebert, E., Rothbächer, U., Pjeta, R., Wunderer, J., Zurovec, M., & Aldred, N. (2021). Omics-based molecular analyses of adhesion by aquatic invertebrates. Biological reviews of the Cambridge Philosophical Society, 96(3), 1051–1075. 10.1111/brv.12691

6. Almeida, M., Reis, R. L., & Silva, T. H. (2020). Marine invertebrates are a source of bioadhesives with biomimetic interest. Materials science & engineering. C, Materials for biological applications, 108, 110467. 10.1016/j.msec.2019.110467

7. Marlétaz, F., Peijnenburg, K. T. C. A., Goto, T., Satoh, N., & Rokhsar, D. S. (2019). A New Spiralian Phylogeny Places the Enigmatic Arrow Worms among Gnathiferans. Current biology: CB, 29(2), 312–318.e3. 10.1016/j.cub.2018.11.042

8. Müller, et al. 2019. Chaetognatha; in: Handbook of Zoology. Ed. Schmidt-Rhaesa.

9. John (1933). Habits, Structure, and Development of Spadella cephaloptera. Quarterly Journal of Microscopical Science 75: 625–96

10. Rieger V, Perez Y, Müller CHG, Lacalli T, Hansson BS, Harzsch S. Development of the nervous system in hatchlings of Spadella cephaloptera (Chaetognatha), and implications for nervous system evolution in Bilateria. Dev Growth Differ. 2011;53:740–59.

11. Arnaud, J., Brunet, M., Casanova, J. P., Mazza, J., & Pasqualini, V. (1996). Morphology and ultrastructure of the gut in Spadella cephaloptera (chaetognatha). Journal of morphology, 228(1), 27–44. 10.1002/(SICI)1097-4687(199604)228:1<27::AID-JMOR3>3.0.CO;2-M

12. Perez, Y., Casanova, J., & Mazza, J. (2000). Changes in the structure and ultrastructure of the intestine of Spadella cephaloptera (Chaetognatha) during feeding and starvation experiments. Journal of experimental marine biology and ecology, 253(1), 1–15. 10.1016/s0022-0981(00)00228-8

13. Ordoñez, J. F., & Wollesen, T. (2024). Unfolding the ventral nerve center of chaetognaths. Neural development, 19(1), 5. 10.1186/s13064-024-00182-6

14. Robertson, H. E., Sebé-Pedrós, A., Saudemont, B., Loe-Mie, Y., Zakrzewski, A. C., Grau-Bové, X., Mailhe, M. P., Schiffer, P., Telford, M. J., & Marlow, H. (2024). Single cell atlas of Xenoturbella bocki highlights limited cell-type complexity. Nature communications, 15(1), 2469. 10.1038/s41467-024-45956-y

15. Hulett, R.E., Kimura, J.O., Bolaños, D.M. et al. Acoel single-cell atlas reveals expression dynamics and heterogeneity of adult pluripotent stem cells. Nat Commun 14, 2612 (2023). 10.1038/s41467-023-38016-4

16. Piovani, L., Leite, D. J., Yañez Guerra, L. A., Simpson, F., Musser, J. M., Salvador-Martínez, I., Marlétaz, F., Jékely, G., & Telford, M. J. (2023). Single-cell atlases of two lophotrochozoan larvae highlight their complex evolutionary histories. Science advances, 9(31), eadg6034. 10.1126/sciadv.adg6034

17. Paganos, P., Voronov, D., Musser, J. M., Arendt, D., & Arnone, M. I. (2021). Single-cell RNA sequencing of the Strongylocentrotus purpuratus larva reveals the blueprint of major cell types and nervous system of a non-chordate deuterostome. eLife, 10, e70416. 10.7554/eLife.70416

18. Sim, S. B., Corpuz, R. L., Simmonds, T. J., & Geib, S. M. (2022). HiFiAdapterFilt, a memory efficient read processing pipeline, prevents occurrence of adapter sequence in PacBio HiFi reads and their negative impacts on genome assembly. BMC genomics, 23(1), 157. 10.1186/s12864-022-08375-1

19. Cheng, H., Concepcion, G.T., Feng, X., Zhang, H., Li H. (2021) Haplotype-resolved de novo assembly using phased assembly graphs with hifiasm. Nat Methods, 18:170–175. 10.1038/s41592-020-01056-5

20. Gurevich, A., Saveliev, V., Vyahhi, N., & Tesler, G. (2013). QUAST: quality assessment tool for genome assemblies. Bioinformatics (Oxford, England), 29(8), 1072–1075. 10.1093/bioinformatics/btt086

21. Simão, F. A., Waterhouse, R. M., Ioannidis, P., Kriventseva, E. V., & Zdobnov, E. M. (2015). BUSCO: assessing genome assembly and annotation completeness with single-copy orthologs. Bioinformatics (Oxford, England), 31(19), 3210–3212. 10.1093/bioinformatics/btv351

22. Wollesen, T., Rodriguez Monje, S. V., Oel, A. P., & Arendt, D. (2023). Characterization of eyes, photoreceptors, and opsins in developmental stages of the arrow worm Spadella cephaloptera (Chaetognatha). Journal of experimental zoology. Part B, Molecular and developmental evolution, 340(5), 342–353. 10.1002/jez.b.23193

23. Flynn, J. M., Hubley, R., Goubert, C., Rosen, J., Clark, A. G., Feschotte, C., & Smit, A. F. (2020). RepeatModeler2 for automated genomic discovery of transposable element families. Proceedings of the National Academy of Sciences of the United States of America, 117(17), 9451–9457. 10.1073/pnas.1921046117

24. Tarailo-Graovac, M., & Chen, N. (2009). Using RepeatMasker to identify repetitive elements in genomic sequences. Current protocols in bioinformatics, Chapter 4, 4.10.1–4.10.14. 10.1002/0471250953.bi0410s25

25. Hoff, K. J., Lange, S., Lomsadze, A., Borodovsky, M., & Stanke, M. (2016). BRAKER1: unsupervised RNA-Seq-based genome annotation with GeneMark-ET and AUGUSTUS. Bioinformatics, 32(5), 767–769.

26. Bruna, T., Hoff, K.J., Lomsadze, A., Stanke, M., & Borodovsky, M. (2021). BRAKER2: Automatic Eukaryotic Genome Annotation with GeneMark-EP+ and AUGUSTUS Supported by a Protein Database. NAR Genomics and Bioinformatics 3(1), lqaa108.

27. Hoff, K. J., Lomsadze, A., Borodovsky, M., & Stanke, M. (2019). Whole-genome annotation with BRAKER. In Gene Prediction (pp. 65–95). Humana, New York, NY.

28. Bruna, T., Lomsadze, A., & Borodovsky, M. (2023). GeneMark-ETP: Automatic Gene Finding in Eukaryotic Genomes in Consistence with Extrinsic Data. bioRxiv, 10.1101/2023.01.13.524024.

29. Buchfink, B., Xie, C., & Huson, D. H. (2015). Fast and sensitive protein alignment using DIAMOND. Nature Methods, 12(1), 59.

30. Gotoh, O. (2008). A space-efficient and accurate method for mapping and aligning cDNA sequences onto genomic sequence. Nucleic acids research, 36(8), 2630–2638.

31. Iwata, H., & Gotoh, O. (2012). Benchmarking spliced alignment programs including Spaln2, an extended version of Spaln that incorporates additional species-specific features. Nucleic acids research, 40(20), e161–e161.

32. Kovaka, S., Zimin, A. V., Pertea, G. M., Razaghi, R., Salzberg, S. L., & Pertea, M. (2019). Transcriptome assembly from long-read RNA-seq alignments with StringTie2. Genome biology, 20(1):1–13.

33. Pertea, G., & Pertea, M. (2020). GFF utilities: GffRead and GffCompare. F1000Research, 9.

34. Gabriel, L., Bruna, T., Hoff, K. J., Ebel, M., Lomsadze, A., Borodovsky, M., & Stanke, M. (2023). BRAKER3: Fully Automated Genome Annotation Using RNA-Seq and Protein Evidence with GeneMark-ETP, AUGUSTUS and TSEBRA. bioRxiv, 10.1101/2023.06.10.544449.

35. Stanke, M., Diekhans, M., Baertsch, R., & Haussler, D. (2008). Using native and syntenically mapped cDNA alignments to improve de novo gene finding. Bioinformatics, 24(5), 637–644.

36. Stanke, M., Schöffmann, O., Morgenstern, B., & Waack, S. (2006). Gene prediction in eukaryotes with a generalized hidden Markov model that uses hints from external sources. BMC Bioinformatics, 7(1), 62.

37. Gabriel, L., Hoff, K. J., Bruna, T., Borodovsky, M., & Stanke, M. (2021). TSEBRA: transcript selector for BRAKER. BMC Bioinformatics, 22:566.

38. Kim, D., Paggi, J. M., Park, C., Bennett, C., & Salzberg, S. L. (2019). Graph-based genome alignment and genotyping with HISAT2 and HISAT-genotype. Nature biotechnology, 37(8), 907–915. 10.1038/s41587-019-0201-4

39. Kuznetsov, D., Tegenfeldt, F., Manni, M., Seppey, M., Berkeley, M., Kriventseva, E. V., & Zdobnov, E. M. (2023). OrthoDB v11: annotation of orthologs in the widest sampling of organismal diversity. Nucleic Acids Research, 51(D1), D445–D451.

40. Li, W., & Godzik, A. (2006). Cd-hit: a fast program for clustering and comparing large sets of protein or nucleotide sequences. Bioinformatics (Oxford, England), 22(13), 1658–1659. 10.1093/bioinformatics/btl158

41. Haas, B. J., Papanicolaou, A., Yassour, M., Grabherr, M., Blood, P. D., Bowden, J., Couger, M. B., Eccles, D., Li, B., Lieber, M., MacManes, M. D., Ott, M., Orvis, J., Pochet, N., Strozzi, F., Weeks, N., Westerman, R., William, T., Dewey, C. N., Henschel, R.,…Regev, A. (2013). De novo transcript sequence reconstruction from RNA-seq using the Trinity platform for reference generation and analysis. Nature protocols, 8(8), 1494–1512. 10.1038/nprot.2013.084

42. Camacho, C., Coulouris, G., Avagyan, V., Ma, N., Papadopoulos, J., Bealer, K., & Madden, T. L. (2009). BLAST+: architecture and applications. BMC bioinformatics, 10, 421. 10.1186/1471-2105-10-421

43. Bairoch, A., & Apweiler, R. (2000). The SWISS-PROT protein sequence database and its supplement TrEMBL in 2000. Nucleic acids research, 28(1), 45–48. 10.1093/nar/28.1.45

44. Eddy S. R. (2011). Accelerated Profile HMM Searches. PLoS computational biology, 7(10), e1002195. 10.1371/journal.pcbi.1002195

45. Hernández-Plaza, A., Szklarczyk, D., Botas, J., Cantalapiedra, C. P., Giner-Lamia, J., Mende, D. R., Kirsch, R., Rattei, T., Letunic, I., Jensen, L. J., Bork, P., von Mering, C., & Huerta-Cepas, J. (2023). eggNOG 6.0: enabling comparative genomics across 12 535 organisms. Nucleic acids research, 51(D1), D389–D394. 10.1093/nar/gkac1022

46. Sarropoulos, I., Sepp, M., Frömel, R., Leiss, K., Trost, N., Leushkin, E., Okonechnikov, K., Joshi, P., Giere, P., Kutscher, L. M., Cardoso-Moreira, M., Pfister, S. M., & Kaessmann, H. (2021). Developmental and evolutionary dynamics of cis-regulatory elements in mouse cerebellar cells. Science (New York, N.Y.), 373(6558), eabg4696. 10.1126/science.abg4696

47. Hao, Y., Stuart, T., Kowalski, M. H., Choudhary, S., Hoffman, P., Hartman, A., Srivastava, A., Molla, G., Madad, S., Fernandez-Granda, C., & Satija, R. (2024). Dictionary learning for integrative, multimodal and scalable single-cell analysis. Nature biotechnology, 42(2), 293– 304. 10.1038/s41587-023-01767-y

48. Baran, Y., Bercovich, A., Sebe-Pedros, A., Lubling, Y., Giladi, A., Chomsky, E., Meir, Z., Hoichman, M., Lifshitz, A., & Tanay, A. (2019). MetaCell: analysis of single-cell RNA-seq data using K-nn graph partitions. Genome biology, 20(1), 206. 10.1186/s13059-019-1812-2

49. Jette, M.A., Wickberg, T. (2023). Architecture of the Slurm Workload Manager. In: Klusáček, D., Corbalán, J., Rodrigo, G.P. (eds) Job Scheduling Strategies for Parallel Processing. JSSPP 2023. Lecture Notes in Computer Science, vol 14283. Springer, Cham. 10.1007/978-3-031-43943-8_1

50. Duruz, J., Kaltenrieder, C., Ladurner, P., Bruggmann, R., Martìnez, P., & Sprecher, S. G. (2021). Acoel Single-Cell Transcriptomics: Cell Type Analysis of a Deep Branching Bilaterian. Molecular biology and evolution, 38(5), 1888–1904. 10.1093/molbev/msaa333

51. Müller, C.H.G., Rieger, V., Perez, Y. et al. Immunohistochemical and ultrastructural studies on ciliary sense organs of arrow worms (Chaetognatha). Zoomorphology 133, 167– 189 (2014). 10.1007/s00435-013-0211-6

52. Xu, X., Xu, M., Zhou, X., Jones, O. B., Moharomd, E., Pan, Y., Yan, G., Anthony, D. D., & Isaacs, W. B. (2013). Specific structure and unique function define the hemicentin. Cell & bioscience, 3(1), 27. 10.1186/2045-3701-3-27

53. Malicki J. (2012). Who drives the ciliary highway?. Bioarchitecture, 2(4), 111–117. 10.4161/bioa.21101

54. Kraut, R., & Zinn, K. (2004). Roundabout 2 regulates migration of sensory neurons by signaling in trans. Current biology: CB, 14(15), 1319–1329. 10.1016/j.cub.2004.07.052

55. Mahen R. (2021). The structure and function of centriolar rootlets. Journal of cell science, 134(16), jcs258544. 10.1242/jcs.258544

56. Vasconcelos, A. C. N., Streit, D. P., Jr, Octavera, A., Miwa, M., Kabeya, N., & Yoshizaki, G. (2019). The germ cell marker dead end reveals alternatively spliced transcripts with dissimilar expression. Scientific reports, 9(1), 2407. 10.1038/s41598-019-39101-9

57. Babin, P. J., Bogerd, J., Kooiman, F. P., Van Marrewijk, W. J., & Van der Horst, D. J. (1999). Apolipophorin II/I, apolipoprotein B, vitellogenin, and microsomal triglyceride transfer protein genes are derived from a common ancestor. Journal of molecular evolution, 49(1), 150–160. 10.1007/pl00006528

58. Sanes J. R. (2003). The basement membrane/basal lamina of skeletal muscle. The Journal of biological chemistry, 278(15), 12601–12604. 10.1074/jbc.R200027200

59. Lv, Y., Pang, X., Cao, Z., Song, C., Liu, B., Wu, W., & Pang, Q. (2024). Evolution and Function of the Notch Signaling Pathway: An Invertebrate Perspective. International journal of molecular sciences, 25(6), 3322. 10.3390/ijms25063322

60. Gao, J., Liu, C. F., Liu, P. P., & Wang, X. W. (2024). Double-stranded RNA induces antiviral transcriptional response through the Dicer-2/Ampk/FoxO axis in an arthropod. Proceedings of the National Academy of Sciences of the United States of America, 121(31), e2409233121. 10.1073/pnas.2409233121

61. Liu, Z., Zhou, Z., Wang, L., Qiu, L., Zhang, H., Wang, H., & Song, L. (2016). CgA1AR-1 acts as an alpha-1 adrenergic receptor in oyster Crassostrea gigas mediating both cellular and humoral immune response. Fish & shellfish immunology, 58, 50–58. 10.1016/j.fsi.2016.09.022

62. Sania, R. E., Cardoso, J. C. R., Louro, B., Marquet, N., & Canário, A. V. M. (2021). A new subfamily of ionotropic glutamate receptors unique to the echinoderms with putative sensory role. Molecular ecology, 30(24), 6642–6658. 10.1111/mec.16206

63. Palazzo, O., Rass, M., & Brembs, B. (2020). Identification of FoxP circuits involved in locomotion and object fixation in Drosophila. Open biology, 10(12), 200295. 10.1098/rsob.200295

64. Lamanna, F., Hervas-Sotomayor, F., Oel, A. P., Jandzik, D., Sobrido-Cameán, D., Santos-Durán, G. N., Martik, M. L., Stundl, J., Green, S. A., Brüning, T., Mößinger, K., Schmidt, J., Schneider, C., Sepp, M., Murat, F., Smith, J. J., Bronner, M. E., Rodicio, M. C., Barreiro-Iglesias, A., Medeiros, D. M.,…Kaessmann, H. (2023). A lamprey neural cell type atlas illuminates the origins of the vertebrate brain. Nature ecology & evolution, 7(10), 1714–1728. 10.1038/s41559-023-02170-1

65. Sur A and Meyer NP (2021) Resolving Transcriptional States and Predicting Lineages in the Annelid *Capitella teleta* Using Single-Cell RNAseq. Front. Ecol. Evol. 8:618007. doi: 10.3389/fevo.2020.618007

66. Borba, C., Kourakis, M. J., Miao, Y., Guduri, B., Deng, J., & Smith, W. C. (2024). Whole Nervous System Expression of Glutamate Receptors Reveals Distinct Receptor Roles in Sensorimotor Circuits. eNeuro, 11(9), ENEURO.0306-24.2024. 10.1523/ENEURO.0306-24.2024

67. Lind, U., Alm Rosenblad, M., Hasselberg Frank, L., Falkbring, S., Brive, L., Laurila, J. M., Pohjanoksa, K., Vuorenpää, A., Kukkonen, J. P., Gunnarsson, L., Scheinin, M., Mårtensson Lindblad, L. G., & Blomberg, A. (2010). Octopamine receptors from the barnacle balanus improvisus are activated by the alpha2-adrenoceptor agonist medetomidine. Molecular pharmacology, 78(2), 237–248. 10.1124/mol.110.063594

68. Xia S, Chiang AS. NMDA Receptors in Drosophila. In: Van Dongen AM, editor. Biology of the NMDA Receptor. Boca Raton (FL): CRC Press/Taylor & Francis; 2009. Chapter 10. Available from: https://www.ncbi.nlm.nih.gov/books/NBK5286/

69. Paganos, P., Voronov, D., Musser, J. M., Arendt, D., & Arnone, M. I. (2021). Single-cell RNA sequencing of the *Strongylocentrotus purpuratus* larva reveals the blueprint of major cell types and nervous system of a non-chordate deuterostome. eLife, 10, e70416. 10.7554/eLife.70416

70. Nishino, A., Okamura, Y., Piscopo, S., & Brown, E. R. (2010). A glycine receptor is involved in the organization of swimming movements in an invertebrate chordate. BMC neuroscience, 11, 6. 10.1186/1471-2202-11-6

71. Ramos-Vicente, D., Grant, S. G., & Bayés, À. (2021). Metazoan evolution and diversity of glutamate receptors and their auxiliary subunits. Neuropharmacology, 195, 108640. 10.1016/j.neuropharm.2021.108640

72. Zeilhofer, H. U., Acuña, M. A., Gingras, J., & Yévenes, G. E. (2018). Glycine receptors and glycine transporters: targets for novel analgesics?. Cellular and molecular life sciences: CMLS, 75(3), 447–465. 10.1007/s00018-017-2622-x

73. Xuereb, B., Lefèvre, E., Garric, J., & Geffard, O. (2009). Acetylcholinesterase activity in Gammarus fossarum (Crustacea Amphipoda): linking AChE inhibition and behavioural alteration. Aquatic toxicology (Amsterdam, Netherlands), 94(2), 114–122. 10.1016/j.aquatox.2009.06.010

74. Horie, T., Nakagawa, M., Sasakura, Y., Kusakabe, T. G., & Tsuda, M. (2010). Simple motor system of the ascidian larva: neuronal complex comprising putative cholinergic and GABAergic/glycinergic neurons. Zoological science, 27(2), 181–190. 10.2108/zsj.27.181

75. Mullen, G. P., Mathews, E. A., Saxena, P., Fields, S. D., McManus, J. R., Moulder, G., Barstead, R. J., Quick, M. W., & Rand, J. B. (2006). The Caenorhabditis elegans snf-11 gene encodes a sodium-dependent GABA transporter required for clearance of synaptic GABA. Molecular biology of the cell, 17(7), 3021–3030. 10.1091/mbc.e06-02-0155

76. Papillon, D., Perez, Y., Fasano, L., Le Parco, Y., & Caubit, X. (2005). Restricted expression of a median Hox gene in the central nervous system of chaetognaths. Development genes and evolution, 215(7), 369–373. 10.1007/s00427-005-0483-z

77. Luparello, C., Mauro, M., Lazzara, V., & Vazzana, M. (2020). Collective Locomotion of Human Cells, Wound Healing and Their Control by Extracts and Isolated Compounds from Marine Invertebrates. Molecules (Basel, Switzerland), 25(11), 2471. 10.3390/molecules25112471

78. Hu, C., Xu, Y., Wen, J., Zhang, L., Fan, S., & Su, T. (2010). Larval development and juvenile growth of the sea cucumber Stichopus sp. (Curry fish). Aquaculture, 300(1–4), 73–79. 10.1016/j.aquaculture.2009.09.033

79. Szulgit G. (2007). The echinoderm collagen fibril: a hero in the connective tissue research of the 1990s. BioEssays: news and reviews in molecular, cellular and developmental biology, 29(7), 645–653. 10.1002/bies.20597

80. Zeng, F., Wunderer, J., Salvenmoser, W., Hess, M. W., Ladurner, P., & Rothbächer, U. (2019). Papillae revisited and the nature of the adhesive secreting collocytes. Developmental biology, 448(2), 183–198. 10.1016/j.ydbio.2018.11.012

81. Davey, P. A., Rodrigues, M., Clarke, J. L., & Aldred, N. (2019). Transcriptional characterisation of the Exaiptasia pallida pedal disc. BMC genomics, 20(1), 581. 10.1186/s12864-019-5917-5

82. Ventura, I., Harman, V., Beynon, R. J., & Santos, R. (2023). Glycoproteins Involved in Sea Urchin Temporary Adhesion. Marine drugs, 21(3), 145. 10.3390/md21030145

83. Kang, V., Lengerer, B., Wattiez, R., & Flammang, P. (2020). Molecular insights into the powerful mucus-based adhesion of limpets (*Patella vulgata* L.). Open biology, 10(6), 200019. 10.1098/rsob.200019

84. Wunderer, J., Lengerer, B., Pjeta, R., Bertemes, P., Kremser, L., Lindner, H., Ederth, T., Hess, M. W., Stock, D., Salvenmoser, W., & Ladurner, P. (2019). A mechanism for temporary bioadhesion. Proceedings of the National Academy of Sciences of the United States of America, 116(10), 4297–4306. 10.1073/pnas.1814230116

